# Extrasynaptic volume transmission: A novel route for neuropeptide signaling in nematodes

**DOI:** 10.1101/2020.08.07.240440

**Authors:** Louise E. Atkinson, Yang Liu, Fiona McKay, Elke Vandewyer, Charles Viau, Allister Irvine, Bruce A. Rosa, Zihui Li, Nikki J. Marks, Aaron G. Maule, Makedonka Mitreva, Isabel Beets, Lingjun Li, Angela Mousley

**Author notes:** Corresponding author **Email:** (AM).

## Abstract

Neural circuit synaptic connectivities (the connectome) provide the anatomical foundation for our understanding of nematode nervous system function. However, other non-synaptic routes of communication are known in invertebrates including extrasynaptic volume transmission (EVT), which enables short- and/or long-range communication in the absence of synaptic connections. Although EVT has been highlighted as a facet of *Caenorhabditis elegans* neurosignaling, no experimental evidence identifies body cavity fluid (pseudocoelomic fluid; PCF) as a vehicle for either neuropeptide or biogenic amine transmission. In the parasitic nematode *Ascaris suum* FMRFamide-like peptides encoded on *flp*-18 potently stimulate female reproductive organs but are only expressed in cells that are anatomically distant from the reproductive organ, with no known synaptic connections to this tissue. Here we report a new non-synaptic mode of signaling in nematodes mediated by neuropeptides within the PCF. Our data show that: (i) *A. suum* PCF (As-PCF) contains a catalogue of neuropeptides including FMRFamide-like peptides and neuropeptide-like proteins; (ii) the *A. suum* FMRFamide-like peptide As-FLP-18A dominates the As-PCF peptidome; (iii) As-PCF potently modulates nematode reproductive muscle function *ex vivo*, mirroring the effects of synthetic FLP-18 peptides; (iv) As-PCF activates the *C. elegans* FLP-18 receptors NPR-4 and -5; (v) As-PCF alters *C. elegans* behavior and, (vi) FLP-18 and FLP-18 receptors display pan-phylum distribution in nematodes. Here we provide the first direct experimental evidence that supports an extrasynaptic volume route for neuropeptide transmission in nematodes. These data demonstrate non-synaptic signaling within the nematode functional connectome and are pertinent to receptor deorphanisation approaches underpinning drug discovery programs for nematode pathogens.

## Introduction

Our current understanding of nematode neuronal circuitry is based on comprehensive *Caenorhabditis elegans* synaptic connectome data [1-3]. The *C. elegans* blueprint underpins fundamental parasitic nematode neurobiology driving anatomical and functional connectomics studies in model parasites such as *Ascaris*, in which the simple neuronal architecture described in *C. elegans* appears to be highly conserved [see 4]. Whilst not experimentally demonstrated in nematodes to date, the existence and significance of additional forms of non-synaptic neuronal communication have been recognized among invertebrates [5, 6]. For example, in crustaceans, extrasynaptic volume transmission (EVT) operates beyond the synaptic connectome, mediating long-range hormonal communication in the absence of neuronal synapses [7, 8]. Notably, EVT has been implicated in nematode neuronal signaling [9-14]. Indeed, the *C. elegans* wired connectome does not always support receptor-ligand interactions that have been functionally linked; there are examples of signaling pathways where receptors are not located co-synaptically with neurons in both monoamine and neuropeptide systems [15-18], supporting a role for EVT in neurotransmission. Further, EVT has been putatively linked to nematode neuropeptide signaling via *in silico* approaches [19]; however, no experimental data are available which test these hypotheses.

Neuropeptides are well known neuromodulators of a wide array of essential neuronal functions in nematodes, including locomotion, reproduction and feeding [see 20 for review]. Despite biological importance, nematode neuropeptide signaling beyond the synaptic connectome is poorly understood. Indeed, the capability and characteristics of neuropeptide EVT in nematodes remain unreported, likely due to the lack of appropriate experimental and analytical tools for the investigation of extracellular communication via body fluid in nematodes including *C. elegans*. The zoonotic gastrointestinal parasite *A. suum* offers a unique opportunity to probe extrasynaptic communication in nematodes. Adult *A. suum*, which are up to 30 cm in length, contain body cavity fluid (pseudocoelomic fluid, As-PCF) that maintains hydrostatic pressure, bathes internal organs and muscle systems, and provides an ideal vehicle for non-neuronal communication. In contrast to *C. elegans*, the size of adult *A. suum* facilitates routine collection of relatively large volumes (∼500 µl/nematode) of As-PCF for bioanalysis. This, in conjunction with the experimental tractability of *Ascaris*, where tissues and organs can be readily dissected for physiology studies [see 20], offers a unique opportunity to investigate EVT in nematodes.

This study exploits peptidomics, nematode physiology, and behavioral bioassays, heterologous expression and bioinformatic approaches to determine the role of nematode As-PCF in EVT of neuropeptides. We show that As-PCF contains a complex array of neuropeptides which modulate nematode behavior. Specifically, we demonstrate that: (i) neuropeptides in As-PCF include FMRFamide-like peptides (FLPs) and neuropeptide-like proteins (NLPs); (ii) As-FLP-18A (GFGDEMSMPGVLRF) is the dominant neuropeptide in As-PCF; (iii) As-PCF is bioactive on the reproductive muscle, altering activity *ex vivo* in a manner similar to that of synthetic As-FLP-18A; (iv) As-PCF activates the *C. elegans* FLP-18 receptors NPR-4 and -5 heterologously expressed in mammalian cells; (v) As-PCF and synthetic As-FLP-18A impact *C. elegans* behavior; and (vi) FLP-18 and the cognate receptors NPR-4 and -5 display pan-distribution across phylum Nematoda. This study provides the first direct experimental evidence that neuropeptide signaling can be mediated non-neuronally by EVT in nematodes. These novel data highlight that non-neuronal, long-range communication is a key facet of neuropeptide signaling in nematodes, adding a new dimension to the interpretation of functional data pertaining to the nematode nervous system.

## Results and discussion

### *Ascaris* PCF contains a rich peptide library

Liquid chromatography coupled with mass spectrometric (LC-MS/MS) analyses of As-PCF (n=24 LC-MS/MS runs representing 67 individual worms; see Supplementary Data S1) reveals that multiple peptide families are present in nematode body cavity fluid. We queried the As-PCF peptidome with *A. suum in silico* peptide libraries (unpublished, constructed in-house) for the major nematode neuropeptide families [FLPs and neuropeptide like proteins (NLPs)], as well as antimicrobial peptides (AMPs). We detected 76 peptides in As-PCF of adult nematodes [41 neuropeptides (6 FLPs, 35 NLPs and 35 AMPs; see Fig 1A and Supplementary Data 1 (S1))]. High-confidence peptide spectrum matches were confirmed for 10 peptides [1% FDR; 21; see Table 1].

**Table 1.**
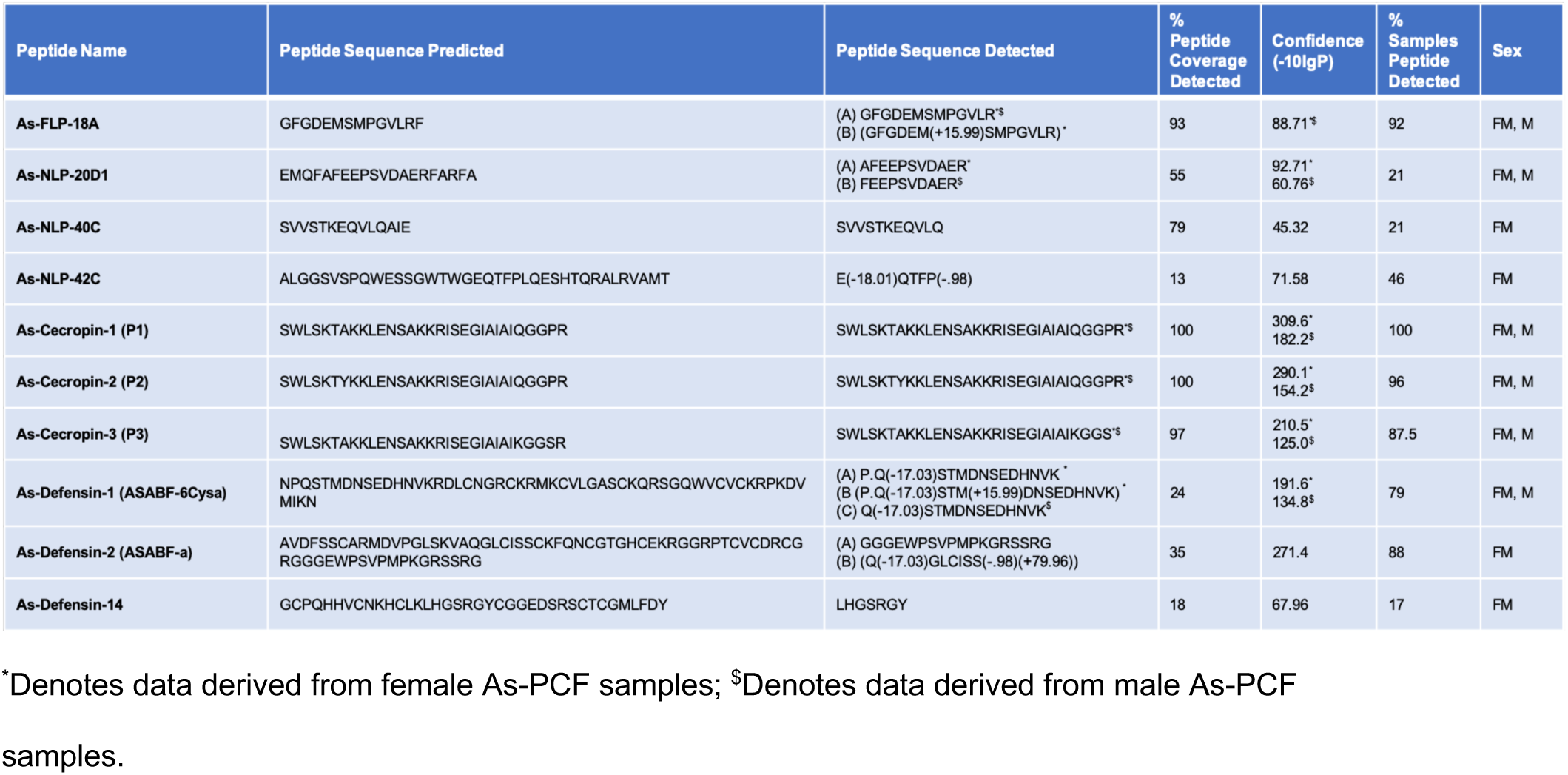
High confidence peptides (>1% FDR) detected by LC-MS/MS in *Ascaris suum* PCF.

**Fig 1.**
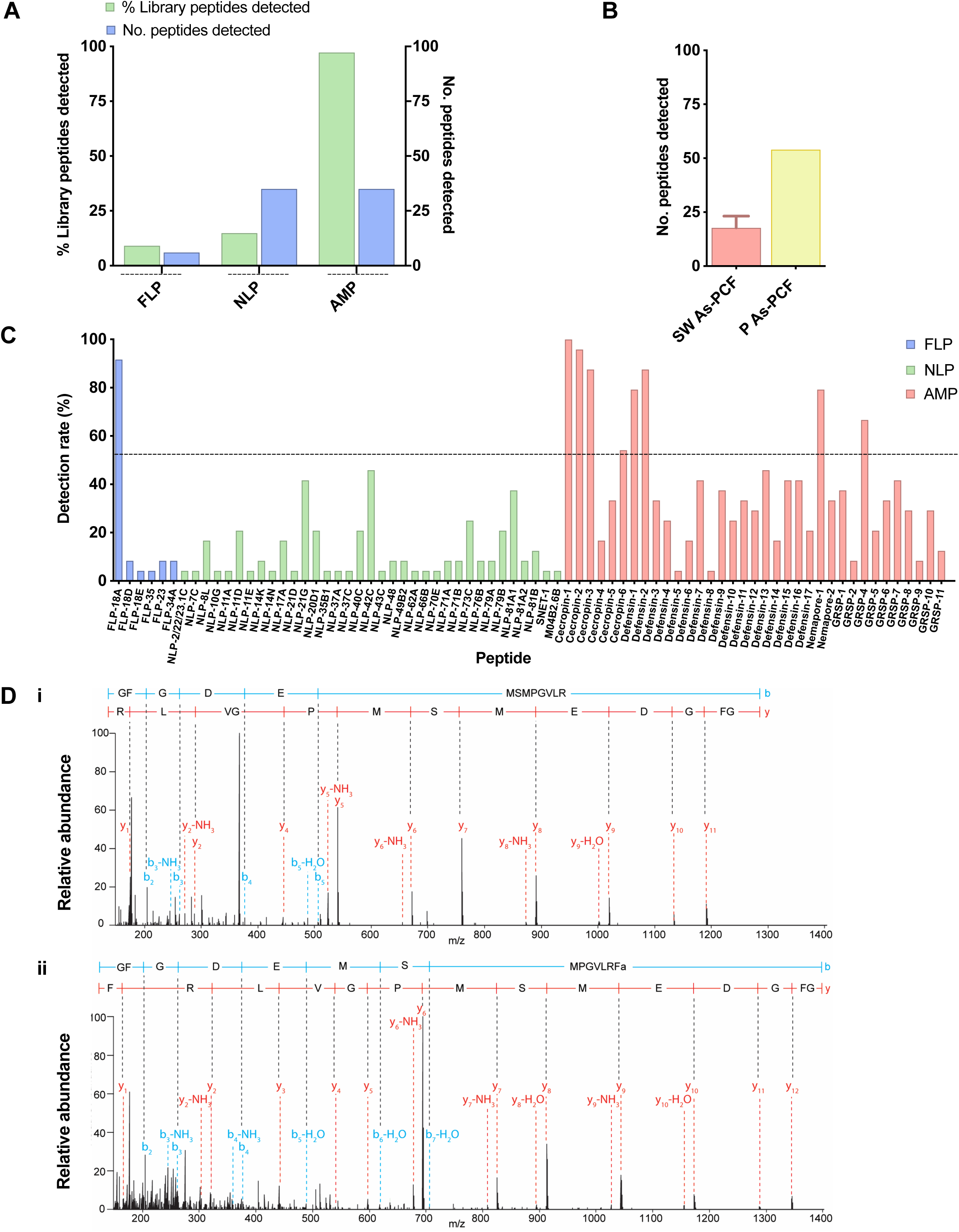
*Ascaris suum* pseudocoelomic fluid (As-PCF) contains a rich neuropeptide library. (A) 41 neuropeptides, including 6 FMRFamide like peptides (FLPs) and 35 neuropeptide like proteins (NLPs), and 35 antimicrobial peptides (AMPs) are present in As-PCF (n=24 LC-MS/MS runs representing 67 individual worms; combined data from single worm, pooled female and male samples). This represents 9.09%, 14.89% and 97.20% of FLP, NLP and AMP peptides (respectively) predicted from the *in silico* libraries used to query the As-PCF LC-MS/MS data. (B) Pooled As-PCF (P As-PCF) derived from female nematodes has a richer neuropeptide complement than single worm As-PCF samples (SW As-PCF) [54 peptides detected in pooled As-PCF (n=17) vs an average of 17.75 ±1.2 peptides across single worm samples (n=20)]. (C) The frequency of detection of individual peptides is variable in As-PCF. Dotted line represents detection rate in 50% of samples. Nine peptides are detected consistently in >50% of As-PCF samples [As-Cecropin P1, As-Cecropin P2, As-FLP-18A, As-Cecropin P3, As-Defensin 2, As-Defensin 1, As-Nemapore 1, As-GRSP-16, As-Cecropin 6]. See Supplementary Data 1 (S1) for all peptide sequences detected. (D) Mass spectra of (i) FLP-18A detected in As-PCF (GFGDEMSMPGVLR) and (ii) As-FLP-18A isotopic standard (GFGDEMSMPGVLRFNH_2_). Amino acid alignments indicate similar fragmentation patterns for As-FLP-18A detected in As-PCF and As-FLP-18A isotopic standard but that As-PCF-derived As-FLP-18A is truncated at the C-terminus. All data are represented as mean ± SEM.

Our experimental approach involved peptidome analyses of single (n=20 worms; 20 LC-MS/MS runs) and pooled adult female (n=17 worms; one LC-MS/MS run) and pooled adult male As-PCF (3 LC-MS/MS runs: n=18, n=6, n=6, worms respectively). This strategy enabled comparison of both pooled and single-worm As-PCF samples, in addition to sex differences. Pooled As-PCF revealed a richer peptide complement than single As-PCF samples; 54 peptides were detected in the pooled sample (n=17 female worms) relative to 17.75 ± 1.2 peptides detected in single worm samples (n=20 worms) (Fig 1B). Six peptides were detected in the pooled sample that were not identified from single worm samples (As-FLP-18D, As-FLP-18E, As-NLP-14N, As-NLP-66B, As-GRSP-2 and As-NLP-14K; see S1). These data indicate that, whilst pooled samples reveal As-PCF peptidome complexity, single worm As-PCF characterization also has utility as an experimental platform for the interrogation of key peptides which are consistently detected in individual worms (see 1% FDR peptides, Table 1 and S1). Male and female As-PCF comparisons revealed two peptides that were male As-PCF specific: As-NLP-43C and a novel FLP, As-FLP-35 (ANTATASWSIEWLMRLNH_2_; see S1). As-FLP-35 is a novel FLP recently predicted from the *A. suum* genome [data unpublished; 22].

Antimicrobial Peptides were the most abundant family of peptides detected in the As-PCF (97% of AMP library predictions); As-Cecropin-1 and 2 [also known as P1 and P2 respectively; 23] were the most highly detected peptides in >96% of samples at >97% coverage (Fig 1A, C and Table 1). FLPs and NLPs displayed greater variability in both the frequency of detection and overall peptide coverage. However, most strikingly, As-FLP-18A equaled the frequency and coverage of the AMPs, dominating the As-PCF peptidome (detected in 92% of samples at >93% coverage; Fig 1A, C and S1). Whilst the number of NLPs detected was greater than the number of FLPs, the majority were low frequency (where approximately half of NLPs were detected only once) with the exception of As-NLP-21G and As-NLP-42C (detected in 42% and 46% of samples, respectively; see Fig 1A-D and S1).

This is the first experimental demonstration of neuropeptides in nematode body cavity fluid. This discovery represents a step-change in the understanding of nematode neuronal signaling and requires that EVT should be considered in the interpretation and comprehension of nematode neurobiology.

### The As-PCF neuropeptidome is compelling in the context of *Caenorhabditis elegans* neurobiology

Neuropeptides are released into the body cavity fluid of invertebrates where they act as neurohormones to regulate many physiological functions [24]. A number of the peptides consistently detected in As-PCF in this study (FLP-18, NLP-21, -37, -40) have been reported in *C. elegans* in cells that link them to the PCF. For example NLP-21, -37 and -40 have been localized to *C. elegans* coelomocytes, which have roles in the endocytosis of secreted proteins from the PCF [11, 12, 25, 26]; this has also been observed for additional neuropeptides, e.g. INS-22 [25], that were not included in the custom peptide library used to probe the As-PCF peptidome in this study. Due to their localization in coelomocyte scavenger cells, it has been postulated that these peptides are released from dense core vesicles and endocytosed. However, until now, their presence in nematode PCF has not been substantiated. In addition, *C. elegans* NLP-37 (PDF-2) has been assigned a long-range neuromodulatory function via release into the PCF as observed for the *Drosophila* homologue [27-30]. The detection of NLP-37 in As-PCF provides evidence for a conserved neuromodulatory role in nematodes. Significantly, a neuromodulatory/neurohormonal role for FLP-18 has also previously been suggested [31]; localization data revealed a physical disconnect between neurons expressing FLP-18 and the location of FLP-18 cognate receptors. The consistent presence of As-FLP-18A in the As-PCF peptidome confirms the potential for FLP-18 peptides to modulate neurobiology via EVT in nematodes.

Based on indirect evidence, a role for EVT of neuropeptides in body cavity fluid has been hypothesised for both *A. suum* and *C. elegans* [9, 14, 19, 31]. However, the data presented here provide the first direct evidence for EVT by neuropeptides in As-PCF. Notably, As-PCF has a restricted peptidome relative to the predicted (*in silico*) whole worm peptidome, consistent with selective release of bioactive peptides into this compartment.

### As-FLP-18A dominates the *Ascaris* PCF neuropeptidome

*As-flp-18* encodes six peptides that share a common C-terminal **PGVLRF-NH**_**2**_ motif (As-FLP-18A, GFGDEMSM**PGVLRF-NH**_**2**_; As-FLP-18B, GM**PGVLRF-NH**_**2**_; As-FLP-18C, AV**PGVLRF-NH**_**2**_; As-FLP-18D, GDV**PGVLRF-NH**_**2**_; As-FLP-18E, SDM**PGVLRF-NH**_**2**_; As-FLP-18F, SM**PGVLRF-NH**_**2**_). As-FLP-18A is the most frequently detected neuropeptide in As-PCF (92% of samples, >93% coverage; Fig 1C), which leads us to hypothesize that As-FLP-18 peptides play an important role in the *A. suum* PCF.

In our LC/MS analysis, we consistently detected a truncated form of As-FLP-18A (GFGDEMSMPGVLR; see Fig 1D) lacking the C-terminal phenylalanine-amide motif required for biological activity (see Fig 2B). Whilst the detection of a truncated peptide could be an artefact of sample processing, we attempted to minimize this possibility as As-PCF was collected and processed rapidly and shipped under temperature-controlled conditions to limit proteolytic degradation. Enhanced detection of truncated As-FLP-18A could be a result of more-ready ionisation of the arginine residue (position -1) in comparison to the intact As-FLP-18A (C-terminal phenylalanine-amide motif). Interestingly, in the pooled As-PCF samples we also detected As-FLP-18D and As-FLP-18E in addition to As-FLP-18A. In these samples, As-FLP-18D and E were also detected in the truncated form without the expected C-terminal phenylalanine-amide motif. Detection of these additional As-FLP peptides in pooled samples, but not in individual worm As-PCF, suggests that these peptides are present at lower concentrations, nearing the LC/MS limit of detection. An alternative hypothesis is that the truncated forms of As-FLP-18A, D and E are produced post-interaction with As-FLP-18 receptors in a signal termination event. Indeed, the E/S products and PCF of nematodes including *A. suum* are known to be peptidase-rich and include a range of cysteine proteases [32-35]. These peptidases may be selectively released at the target receptor to spatiotemporally control peptide interactions and mediate neuropeptide action [36]. In this scenario, peptidase action could manifest in an elevation, and thus an enhanced detection, of truncated versus intact As-FLP-18 peptides. This is consistent with our observations and would contribute to the plasticity of nematode neuronal signaling via EVT. The biological reason for the predominance of As-FLP-18A over the additional five peptides encoded by *As-flp-18* is unclear. Indeed, *As-flp-18* is not predicted to be differentially spliced in *A. suum* as revealed by both genome-level analyses and molecular (PCR) investigation (data not shown). Regardless, truncated As-FLP-18A (synthetic) lacks biological activity in *A. suum* muscle physiology assays in contrast to intact As-FLP-18A (synthetic) and As-PCF, which both induce characteristic FLP-18 responses (see subsequent discussion and Fig 2A and B). These data strongly suggest that both the intact and truncated form of As-FLP-18A are present in As-PCF.

**Fig 2.**
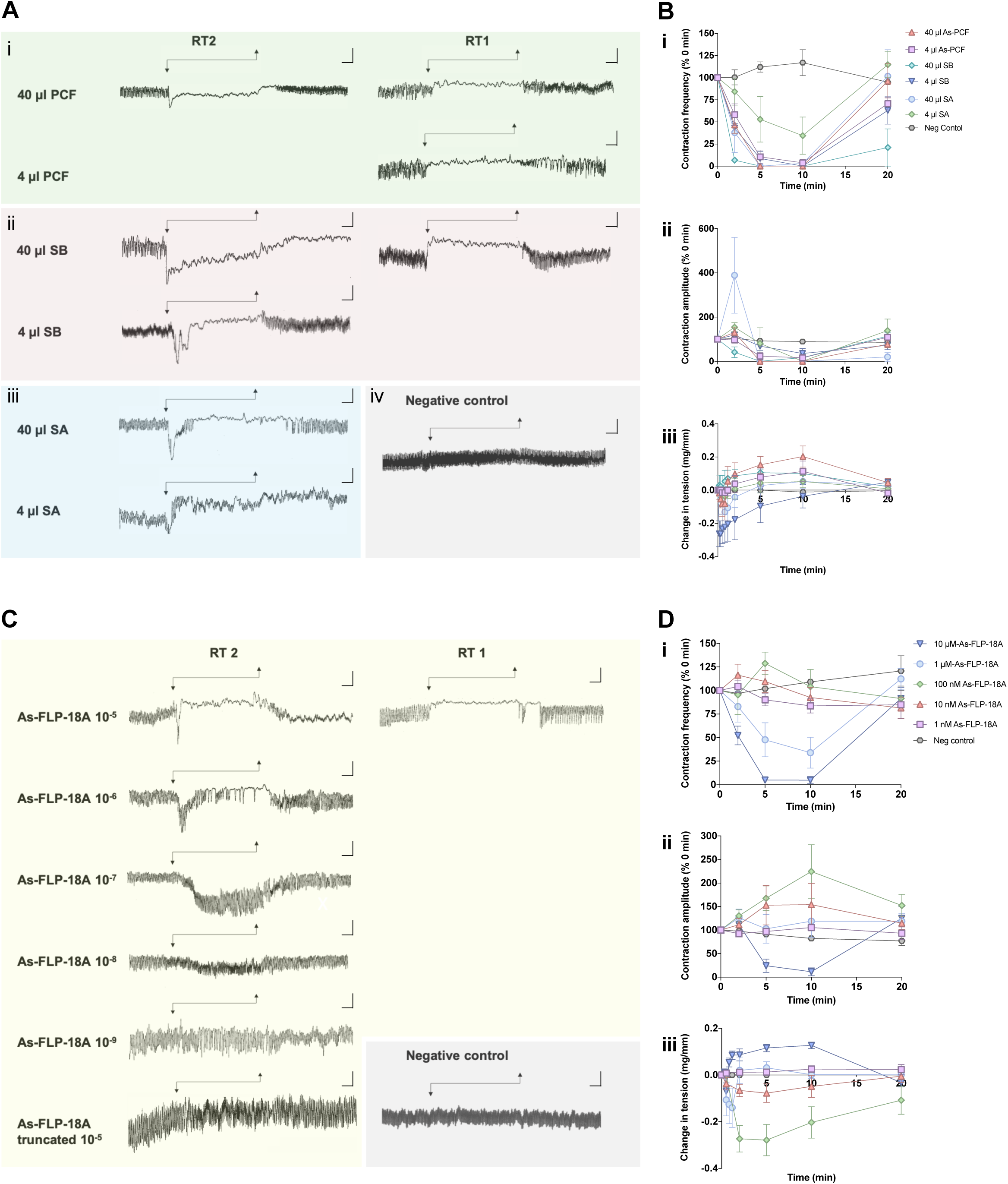
*Ascaris suum* pseudocoelomic fluid (As-PCF) is bioactive on *Ascaris* ovijector tissue and mirrors the synthetic As-FLP-18A peptide response. (A) Representative muscle tension recordings showing the effects of As-PCF on *A. suum* ovijector tissue: (i) As-PCF induces two distinct, concentration dependent, myoactivity profiles (known as response types; RT) on the *A. suum* ovijector; 40 µl As-PCF (equivalent to ≥0.15X As-PCF where 1X is representative of the biological sample) induces either a transient muscle contraction followed by muscle paralysis (50% of preparations; RT-2), or immediate muscle relaxation (50% of preparations; RT-1). 4 µl As-PCF (equivalent to 0.015X As-PCF where 1X is representative of the biological sample) induces an RT-1 response only; (ii) Positive controls [phosphate buffered saline spiked with synthetic As-FLP-18A (GFGDEMSMPGVLRFNH_2_) before C18 peptide purification; SB] induce two distinct, concentration dependent, myoactivity profiles on the *A. suum* ovijector. 40 µl SB (equivalent to final concentration of 1 µM synthetic As-FLP-18A) induces either a RT-2 (50% of preparations), or RT-1 (50% of preparations) response. 4 µl SB (equivalent to final concentration of 0.1 µM synthetic As-FLP-18A) induces an inhibitory RT-1 response only; (iii) Positive controls [phosphate buffered saline spiked with synthetic As-FLP-18A (GFGDEMSMPGVLRFNH_2_) after C18 peptide purification (SA) also induce a concentration dependent myoactivity profile on the *A. suum* ovijector. 40 µl and 4 µl SA controls (equivalent to final concentration of 1 µM and 100 nM synthetic As-FLP-18A, respectively) induce an RT-2 response in 100% of preparations. 40 µl and 4 µl SA controls (equivalent to final concentration of 1 µM and 100 nM synthetic As-FLP-18A, respectively) induce an RT-2 response in 100% of preparations; (iv) Negative control (perfused activation solution only resuspended in ddH_2_O) does not modulate intrinsic ovijector contractility. Test compounds (As-PCF, SB, SA. negative control), were present during the period indicated by arrows. Vertical scales represent 2 mg and horizontal scales represent 2 min. (B) The effects of As-PCF and positive controls (SB and SA) on: (i) contraction frequency, (ii) contraction amplitude and, (iii) muscle tension of the *A. suum* ovijector. Test compounds (As-PCF, SB or SA) were added at 0 min and washed out at 10 min. Data are presented as mean ± SEM [see Supplementary Table 1 (S1 Table)]. (C) Representative muscle tension recordings showing the effects of synthetic As-FLP-18A on *A. suum* ovijector tissue. 1 nM-10 µM synthetic As-FLP-18A induce concentration dependent myoexcitatory effects; 10 µM synthetic As-FLP-18A induces two distinct myoexcitatory profiles consisting with RT-2 (35.7% of preparations), or RT-1 (64.2% of preparations); 1 µM-10 nM synthetic As-FLP-18A induce RT-2 responses only. The truncated form of synthetic As-FLP-18A (GFGDEMSMPGVLR; 10 µM) does not modulate ovijector contractility. Negative control (ddH_2_O) does not modulate intrinsic ovijector contractility. Peptide was present during the period indicated by the arrows. Vertical scales represent 2 mg and horizontal scales represent 2 min. (D) The concentration dependent effects of 1 nM-10 µM synthetic As-FLP-18A on (i) contraction frequency, (ii) contraction amplitude and, (iii) muscle tension of the *A. suum* ovijector. Peptide was added at 0 min and washed out at 10 min. All data are presented as mean ± SEM [see Supplementary Table 2 (S2 Table)].

### The bioactivity of *Ascaris* PCF on *Ascaris* reproductive tissue mirrors that of synthetic As-FLP-18A

Neuropeptides potently modulate the intrinsic muscle activity of *Ascaris* reproductive organs [ovijector; 37, 38-42] defined by five distinct response types (RT-1-RT5), ranging from inhibition of muscle contraction (flaccid paralysis, e.g., RT-1) to increased contraction frequency [e.g. RT5; 38]. As-PCF also potently modulates *Ascaris* reproductive muscle activity. Exogenously applied As-PCF induces two distinct myoactivity profiles on the ovijector that are characteristic of either an RT-1- or RT-2 like response [38; see Figure 2B]. At low concentrations [0.015X As-PCF (where 1X As-PCF represents the native biological sample)], As-PCF consistently inhibits muscle activity in an RT-1 like response [Fig 2A; Supplementary Table 1 (S1 Table)]. At higher concentrations (≥0.15X As-PCF), the response profile included both RT-1 (inhibitory; 50% of preparations) and RT-2 like responses (excitatory; 50% of preparations) (see Fig 2A; S1 Table). All effects were reversible upon As-PCF washout.

Interestingly, in this study we present myoactivity profiles for As-PCF that are consistent with the myoactivity profile of synthetic As-FLP-18A which is also the dominant As-PCF peptide. Synthetic As-FLP-18A induces consistent excitatory effects (10 nM-1 µM) that include transient muscle contraction followed by muscle paralysis [1 µM; RT-2 like; Fig 2B; Supplementary Table 2 (S2 Table)] [38]. At high concentrations (10 µM) synthetic As-FLP-18A induces two distinct myoexcitatory profiles that resemble those observed with high concentrations of As-PCF (RT-1 like, 64.2% of preparations; RT-2 like, 35.7% of preparations; see Fig 2B and S2 Table). These responses were corroborated in synthetic peptide positive controls subjected to the same experimental manipulations as As-PCF (Fig 2B; S2 Table).

As-FLP-18 peptides (PGVLRFNH_2_) have been implicated in locomotion, feeding and reproductive function via whole worm, pharyngeal and ovijector tissue bioassays [see 20]. However, although *As-flp-18* encoded peptides are potent stimulators of the *Ascaris* ovijector, their expression pattern is limited to neurons that are anatomically distinct from, and have no known synaptic connections to, reproductive tissue [as demonstrated by in situ hybridization and MS/MS; 43]. The ovijector receives input from two parallel neuronal cell bodies situated in close proximity to the body wall and the ventral nerve cord (similar positions to *C. elegans* VC4 and VC5) [37]. In this study, RNAseq analysis was conducted on two distinct tissue types: (i) the gonopore region, an area of body wall tissue which contains the neuronal cell bodies which innervate the ovijector and; (ii) the ovijector itself which expresses the post-synaptic receptors. Our tissue-specific differential expression analyses of *As-flp*-18 and the As-FLP-18 cognate GPCR sequelogs *As-npr*-4 and *As-npr*-5 [13, 31, 44], revealed that the neuronal cell bodies innervating the ovijector do not express *As-flp*-18 despite the presence of both cognate receptors in the ovijector tissue (unpublished data manuscript in preparation). These data support the hypothesis that As-PCF-circulating neuropeptides modulate *A. suum* reproductive function. Whilst our data point towards a significant role for As-FLP-18A in modulating reproductive function via EVT, the contribution of other peptides detected in the As-PCF will require further investigation to unravel EVT dynamics.

### *Ascaris* PCF activates heterologously expressed *Caenorhabditis elegans* NPR-4 and NPR-5

To characterize the interaction of As-PCF with the cognate FLP-18 receptors NPR-4 and -5, and to provide further evidence for As-FLP-18 signaling via EVT, we measured the response of Ce-NPR-4 and -5 (expressed in CHO cells) to As-PCF and synthetic As-FLP-18 peptides. Note that attempts to express As-NPR-4 and -5 in the heterologous CHO system were unsuccessful in this study [consistent with difficulties in parasitic nematode GPCR heterologous expression; see 20 for review], requiring the use of the *C. elegans* receptors as surrogates. As-PCF and synthetic As-FLP-18 peptides potently activated Ce-NPR-4 and -5 (synthetic Ce-FLP-18 peptides also elicited the expected responses; see Figs 3A-C). Synthetic As-FLP-18A-E elicited significant Ca^2+^ responses in cells transfected with *C. elegans* NPR-4 and cells transfected with NPR-5, compared to the negative control (cell medium without peptide; see Fig 3A). Synthetic As-FLP-18 peptides did not evoke significant Ca^2+^ responses in CHO cells transfected with an empty vector, confirming that Ce-NPR-4 and -5 mediate the observed physiological responses (Fig 3A). As-PCF also evoked a significant Ca^2+^ response in Ce-NPR-4 or Ce-NPR-5 transfected cells compared to negative controls (see Figs 3B and C). A low-level Ca^2+^ response was observed in cells transfected with an empty control vector as well, indicating that some component(s) of As-PCF interact with receptors endogenously expressed in CHO cells (see Figs 3B and C). Despite this, As-PCF elicited a significantly higher response in cells expressing Ce-NPR-4 or -5 than the negative control (empty vector transfected CHO cells), indicating that a significant component of the As-PCF response is specific to Ce-NPR-4 and -5 activation (see Fig 3B and C respectively).

**Fig 3.**
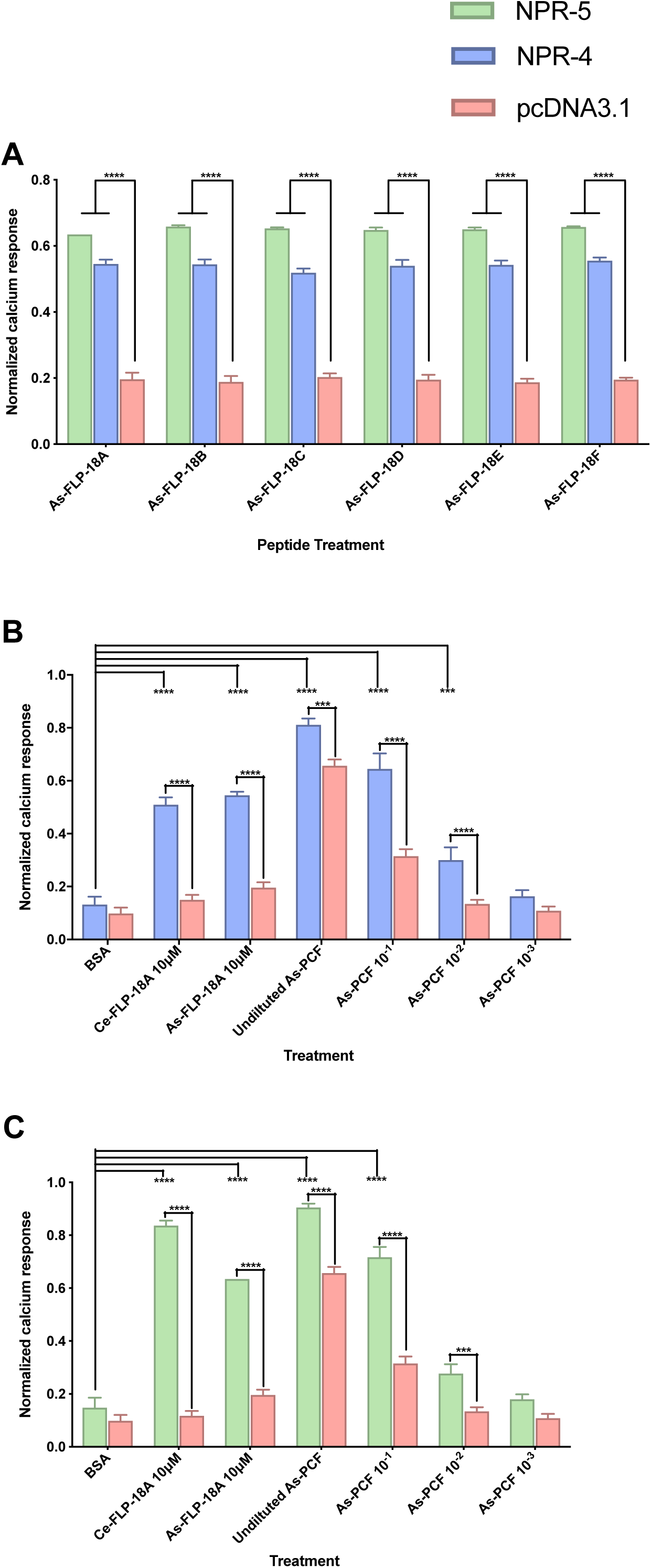
*Ascaris* PCF activates heterologously expressed *Caenorhabditis elegans* neuropeptide receptors NPR-4 and NPR-5 in mammalian cell lines. (A) *C. elegans* NPR-4 (Ce-NPR-4) and -5 (Ce-NPR-5) expressed in CHO cells are activated by synthetic As-FLP-18 peptides (10 μM; As-FLP-18A: GFGDEMSMPGVLRFNH_2_; As-FLP-18B: GMPGVLRFNH_2_; As-FLP-18C: AVPGVLRFNH_2_; As-FLP-18D: GDVPGVLRFNH_2_; As-FLP-18E: SDMPGVLRFNH_2_; As-FLP-18F: SMPGVLRFNH_2_) compared to controls transfected with an empty vector (pcDNA3.1). (B) Ce-NPR-4 is activated by As-PCF in a concentration dependent manner compared to both peptide-free cell medium (BSA) and empty vector (pcDNA3.1) negative controls. The cognate Ce-NPR-4 peptide Ce-FLP-18A (DFDGAMPGVLRFNH_2_) and As-FLP-18A were used as positive controls. (C) Ce-NPR-5 is also activated by As-PCF in a concentration dependent manner compared to both peptide-free cell medium (BSA) and empty vector (pcDNA3.1) negative controls. The cognate Ce-NPR-5 peptide Ce-FLP-18A and As-FLP-18A were used as positive controls. In all cases data are shown as the ratio of peptide/As-PCF response to total calcium response. Error bars represent mean ± SEM., n>3 replicates. Statistical significance of peptide or As-PCF-evoked responses compared with BSA or empty vector controls was determined by two-way ANOVA and Tukey’s multiple comparisons test. *P*-values are denoted by: ***<0.001 and ****<0.0001.

The activation of heterologously expressed Ce-NPR-4 and -5 by As-PCF and synthetic As-FLP-18 peptides provides data that verifies conservation of the FLP-18 interaction with NPR-4 and -5 [31, 44] in *Ascaris* and supports our hypothesis that As-PCF peptides interact with putative FLP-18 receptors *in vivo* via EVT.

### *Ascaris* PCF impacts *Caenorhabditis elegans* growth, reproduction and locomotion

To further investigate the biological activity of As-PCF in nematodes we examined the effects of As-PCF and synthetic As-FLP-18A in *C. elegans* using three behavioral assays which assess nematode growth (body length), reproduction (progeny) and motility (locomotory activity) [45, 46] in *Ce-acs*-20 [cuticle defective mutant to facilitate peptide uptake; 47]. *Caenorhabditis elegans* growth was potently inhibited in a dose-dependent manner by 2% and 5% As-PCF; this effect was mirrored by treatment of *C. elegans* with the dominant As-PCF neuropeptide As-FLP-18A (100 μM, synthetic peptide) (see Fig 4A). In addition, As-PCF dose-dependent effects on reproduction were also evident, where progeny was reduced in the presence of 1, 2 and 5% As-PCF; these effects were also evident in nematodes treated with 100 μM synthetic As-FLP-18A (see Fig 4B). *Caenorhabditis elegans* motility was significantly impaired in the presence of 5, 7.5 and 10% As-PCF as measured by the wMicroTracker (InVivo Biosystems, Oregon, USA); synthetic As-FLP-18A (10 μM) also produced significant inhibition of locomotion (see Fig 4C). In all assays the As-PCF induced modulation of *C. elegans* behavior was consistent with that observed for the synthetic As-FLP-18A peptide (see Fig 4A-C) and corresponds to published literature which demonstrates the modulatory activity of FLP-18 in a number of *C. elegans* behaviors and biological processes [see 48 for review].

**Fig 4.**
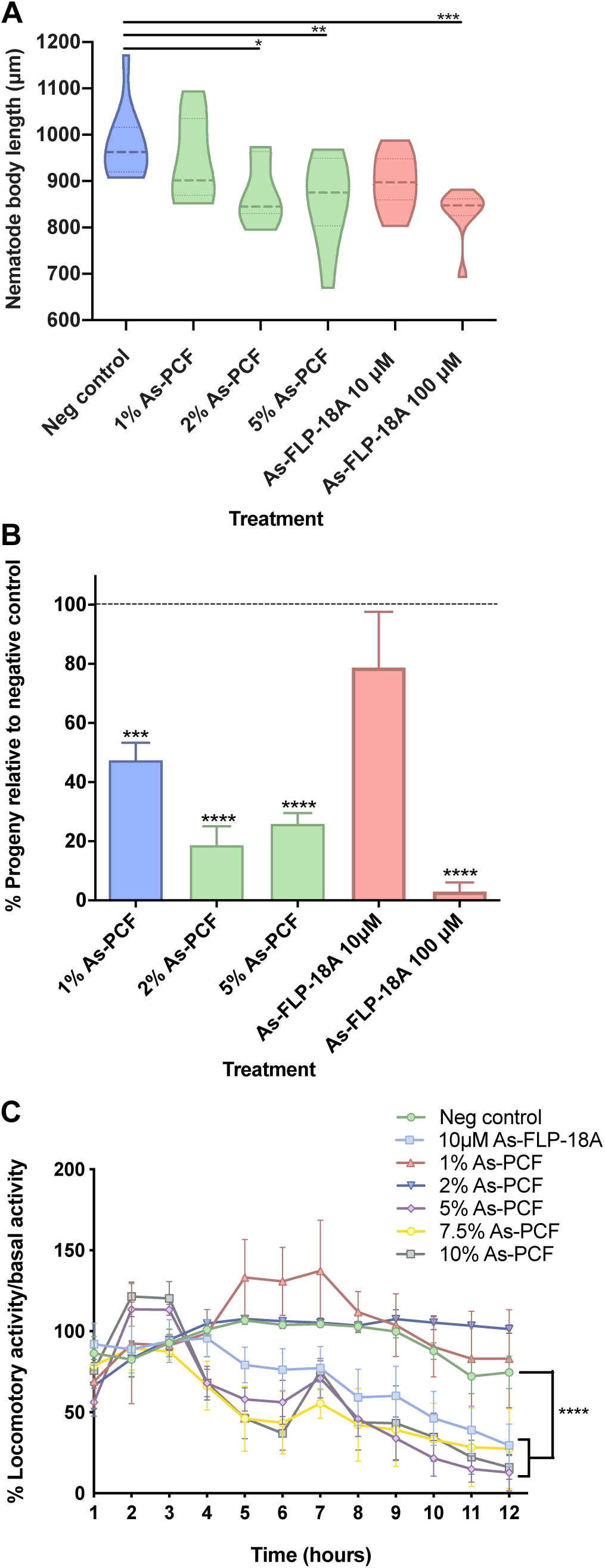
Exogenous application of *Ascaris suum* PCF (As-PCF) and synthetic As-FLP-18A influences *Caenorhabditis elegans* growth, reproduction and motility. (A) *C. elegans* growth (as measured by changes in body length) is significantly reduced in nematodes exposed to 2% and 5% As-PCF and 100 µM synthetic As-FLP-18A (GFGDEMSMPGVLRFNH_2_) compared to the negative control (nematodes exposed to S-medium only). (B) *C. elegans* reproduction (as measured by progeny) is significantly reduced in nematodes exposed to 1%, 2%, 5% As-PCF and 100 µM synthetic As-FLP-18A relative to the negative control (nematodes treated with S-medium only). (C) *C. elegans* motility (as measured locomotory activity over time) is significantly reduced in nematodes exposed to 5%, 7.5%, 10% As-PCF and 10 µM synthetic As-FLP-18A compared to the negative control (nematodes exposed to M9 only). Statistical significance determined by one-way ANOVA and Dunnett’s multiple comparisons test (A and B) or two-way ANOVA and Tukey’s multiple comparisons test (C). *P*-values are denoted by: * <0.05, **<0.01, ***<0.001 and ****<0.0001.

### *flp*-18 and the cognate receptors, *npr*-4 and *npr*-5, display pan-phylum conservation in nematodes

*In silico* analyses of 134 available nematode genomes, representing 109 species, 7 Clades and 3 distinct lifestyles [free-living, animal parasitic and plant parasitic; 49, 50, 51], demonstrates striking *flp*-*18* pan-phylum conservation in 99% of nematode species [see Fig 5 Supplementary Data 2 (S2)]. *flp*-18 is the only neuropeptide gene which displays this level of conservation in phylum Nematoda (unpublished data). These data drive the hypothesis that FLP-18 peptides are fundamental to nematode biology and support the potential for FLP-18 signaling via EVT to be a pan-phylum trait.

**Fig 5.**
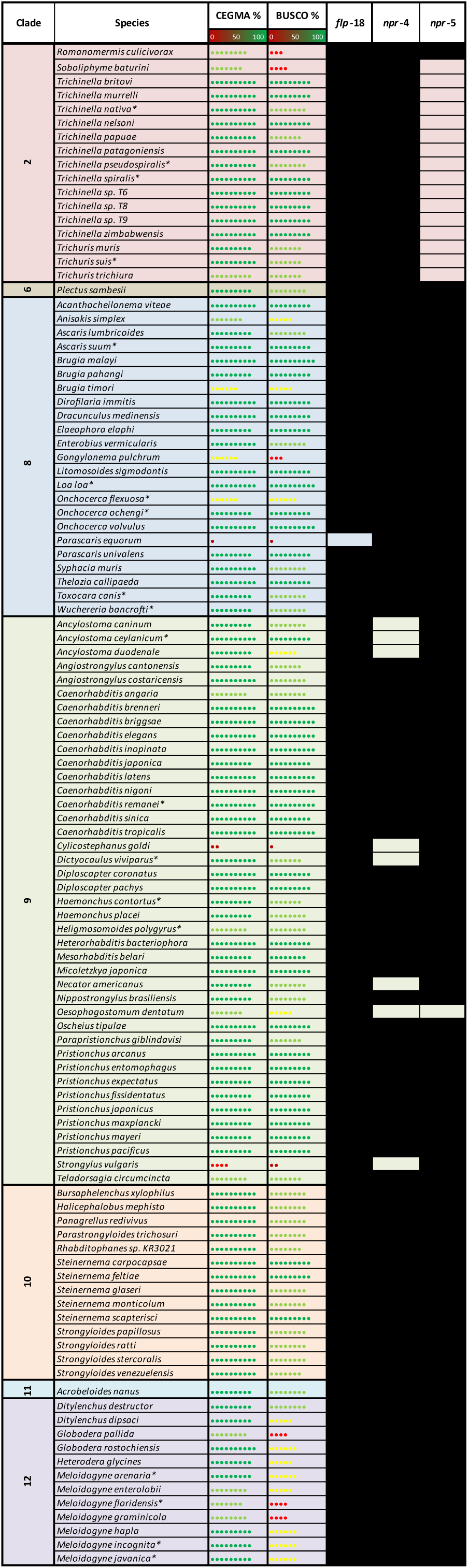
*Caenorhabditis elegans flp*-18, and the cognate FLP-GPCR encoding gene homologues *npr*-4 and *npr*-5 display pan-phylum conservation in nematodes. Pan-phylum HMM analysis of 134 nematode genomes (109 species) demonstrates that *C. elegans flp*-18, *npr-4* and *npr-5* encoding gene homologues are highly conserved (99%, 93%, 84% respectively) in nematodes. Black boxes indicate the presence of a putative gene homologue. Presence/absence identified based on motif conservation and reciprocal BLAST. Nematode species are arranged based on 12 clade designation [58]. Asterisk denotes that multiple genomes were mined per species. Genome quality is indicated by CEGMA and BUSCO scores obtained from Wormbase Parasite version 14 [49-51, 59, 60]. A list of genomes mined, HMM query sequences returned and retrieved gene homologues are detailed in Supplementary Data 2 (S2).

Broad conservation was also noted for the putative FLP-18 receptor encoding genes (*npr*-4 in 93% of species and *npr*-5 in 84% of species; see Fig 5), highlighting their importance across phylum Nematoda. It is interesting to note that, for most of the species examined (98%), the presence of the FLP-18-encoding gene was indicative of the presence of either putative cognate receptor or both and *vice versa* (Fig 5). These data suggest that the FLP-18-NPR-4/5 interaction, which has been functionally validated in *C. elegans, Brugia malayi* [31, 44] and, in this study, in *A. suum*, is highly conserved across nematodes; consequently these receptors hold significant appeal as broad-spectrum drug targets in parasitic nematodes.

## Conclusions

This study reveals a rich neuropeptide profile in the body cavity fluid of a parasitic nematode and, for the first time, provides direct evidence that EVT is an integral part of the nematode functional connectome. These data demonstrate that nematode signaling systems operate within a connectome comprising both wired and wireless components, adding complexity and plasticity to nematode communication frameworks. The biological significance and exploitation of the nematode connectome is significantly enhanced by the wireless data presented here. The combined network infrastructure revealed the experimental tractability of *A. suum* in this study provides a unique and powerful tool to unravel the intricacies of information flow within the nervous system of nematodes and higher organisms.

## Materials and methods

### Collection and maintenance of *Ascaris suum*

Adult *A. suum* were collected at a local abattoir (Karro Food Group Ltd, Cookstown, Northern Ireland), transported to the laboratory in saline (0.9% NaCl), and were maintained in *Ascaris* Ringers Solution (ARS: 13.14 mM NaCl, 9.47 mM CaCl_2_, 7.83 mM MgCl_2_, 12.09 mM C_4_H_11_NO_3_/Tris, 99.96 mM NaC_2_H_3_O_2_,19.64 mM KCl, pH 7.8) at 37 °C until use.

### Maintenance of *Caenorhabditis elegans*

The *C. elegans* cuticle defective mutant strain *acs*-20 [tm3232; 47] was obtained from the National BioResource Project of the Tokyo Women’s Medical University (Tokyo, Japan). *Ce-acs*-20 have defects in the cuticle barrier which increase permeability to small molecules [47]. *Ce-acs-*20 were maintained on Nematode Growth Media (NGM) plates seeded with *Escherichia coli* OP50 (CGC) and synchronized using established methods [52].

### *Ascaris suum* pseudocoelomic fluid collection

As-PCF collection from adult *A. suum* was carried out within 3 hrs of parasite collection. Female worms >20 cm were used in experiments. Between 400 and 1000 µl of As-PCF was collected from each female worm and between 100 and 500 µl of As-PCF was collected from each male worm. As-PCF was obtained from blot-dried (Kimwipes, Fisher Scientific, UK) female worms by holding the specimen vertically above a 2 ml low binding microcentrifuge tube (Fisher Scientific, UK) and carefully snipping <3 mm tissue from the tail (posterior to the anal pore and the major ganglia) using a pair of sterile scissors. AS-PCF was discharged from the worm into the collection tube with the aid of internal turgor pressure and gentle manual compression. Male As-PCF was collected by drying the cuticle and making a 1 cm incision with a sharp scalpel approximately 5 cm posterior to the head (mid body region) and allowing turgor pressure to discharge As-PCF into a 2 ml collection tube; male worms are not amenable to female worm As-PCF collection processes due to the close proximity of male reproductive organs at the posterior tip of the tail which prevents access to the pseudocoelomic cavity. As-PCF was placed on ice immediately after collection. Single worm As-PCF (n=24) and pooled As-PCF analyses (n=17 worms pooled) were conducted for female worms. Male single worm As-PCF analysis was not possible due to the low volume of As-PCF retrieved from individual male worms; As-PCF collected from 30 males was used in pooled analyses

### LC-MS/MS

#### Peptide extraction

Peptide extraction was carried out in two stages. Phase one was conducted at Queen’s University Belfast immediately following As-PCF collection; phase two was conducted at the University of Wisconsin-Madison, WI, USA. *Phase one*: As-PCF was spiked with an equal volume of acidified methanol (90% MeOH, 9% ddH_2_O, 1% glacial CH_3_COOH) immediately following collection (to extract peptides and precipitate large proteins), vortexed and centrifuged for 15 min at 20,000 *g*. The supernatant was retained in a 2 ml low binding microcentrifuge tube (Fisher Scientific, UK). The pellet was resuspended in 500 µl acidified methanol using a sterile plastic pestle. This was repeated twice more and the supernatant from each extraction was retained as above in addition to the final pellet. 250 µl ddH_2_O was added to the supernatant solutions from extractions 2 and 3 to decrease MeOH concentration to 60%. All three supernatant solutions and the final pelleted solids remaining after the third extraction were snap frozen in liquid N_2_ and stored at −80 °C prior to shipping. *Phase two*: upon receipt (University of Wisconsin, Madison), frozen supernatant was thawed and loaded immediately onto a 10 kDa molecular weight cut-off membrane (Sigma-Aldrich, St. Louis, MO) which was rinsed with 0.1 M NaOH and 50/50 MeOH/H_2_O, (v/v) prior to sample loading. The tube holding the membrane was centrifuged at 14,000 *g* for 10 min and the filtrate was dried in a SpeedVac concentrator (Thermo Fisher Scientific, Waltham, MA, USA). The purified sample was resuspended in 150 μl 0.1% formic acid (FA) in H_2_O, desalted by OMIX C18 pipette tips (Agilent Technologies, Santa Clara, CA) and finally eluted into 0.1% FA in 50/50 ACN/H_2_O (v/v). The eluate was concentrated until dry and stored at −80 °C. *LC-MS/MS analysis:* Stored samples were dissolved in 10 μl 0.1% formic acid (FA). Nano LC-MS/MS/MS analysis was carried out using a Waters Nano-Acquity Ultra Performance LC system (Waters Corp, Milford, MA, USA) coupled to a Q-Exactive Quadrupole-Orbitrap mass spectrometer (Thermo Scientific Bremen, Germany). A self-packed column (150 mm length of 1.7 μm C18 with a 3 μm C18 cap) was used for chromatographic separation. The mobile phase involved in online separation was: A, 0.1% FA in H_2_O and B, 0.1% FA in ACN. A 120-min gradient was applied at flow rate of 0.3 μl/min, starting from 100% A. Mobile phase B increased to 10% in 1 min, 35% at 90 min, 95% at 92 min. The gradient remained at 5% A for 10 min, then recovered to 100% A at 105 min. Data were collected under positive electrospray ionization data-dependent mode (DDA), with the top 15 abundant precursor ions selected for HCD fragmentation with listed settings: full-MS, resolution, 70,000; automatic gain control (AGC), 1e6; scan range, m/z 200-2000; dd-MS2, resolution 17,500; AGC, 2e5; isolation window, m/z 2.0; fixed first mass, m/z 100.0; normalized collision energy, 30. Other typical MS parameters were: spray voltage, 2.1 kV; no sheath and auxiliary gas flow; heated capillary temperature, 275 °C. *MS data analysis:* Parent mass error tolerance was 10 ppm and fragment mass error tolerance was 0.02 Da. C-terminal amidation, methionine oxidation and pyro-glutamation were specified as variable post-translational modification (PTMs). No enzyme cleavage was selected within the de novo sequencing. A custom neuropeptide database representing predicted *A. suum* FLPs, NLPs and AMPs was used in the search process. A threshold of false discovery rate (FDR) of 1% was used for data filtration and validation. NanoLC-ESI-MS raw data were analyzed by PEAKS Studio 7 (Bioinformatics Solution Inc., Waterloo, ON, Canada).

### *Ascaris suum* ovijector bioassays

#### Tissue excision and data collection

Adult female *A. suum* ovijector tissue and data collection was carried out as previously described [41]. Briefly, freshly excised *A. suum* ovijector tissue was attached to the recording apparatus and synthetic peptide or As-PCF were added directly to the chamber following tissue equilibration. Muscle activity was recorded for 10 min. Media in the recording chamber was then replaced with fresh HBSS (37°C; Sigma-Aldrich), and the muscle allowed to recover for a further 10 min. If regular baseline activity was achieved following washout further experiments with the specimen were conducted. *FLP-18 bioassays: A. suum* synthetic FLP-18 peptide [As-FLP-18A, GFGDEMSMPGVLRFNH_2_; Genosphere Biotechnologies (France)] was tested for bioactivity on ovijector tissue. Peptide stock solutions (10 mM – 1 µM) were prepared using ddH_2_O such that 4 µl peptide stock solution was added to 4 ml HBSS in the recording chamber to achieve the desired final peptide concentration (10 – 0.001 µM). ddH_2_O was used as a negative control. *As-PCF bioassays:* 3 ml fresh As-PCF was collected from female worms (as described above) and passed through a Sep-Pak^®^ Classic C18 cartridge (Waters, Ireland). Sep-Pak^®^ Classic C18 cartridges were activated with 3 ml activation solution (3 ml Ultrapure Acetonitrile, 0.1 % HPLC grade trifluoroacetic acid) and washed with 6 ml wash solution (6 ml ddH_2_O, 0.1 % HPLC grade trifluoroacetic acid). Following initial flow through, As-PCF effluent was passed through the C18 cartridge twice more to ensure optimal peptide binding. C18 cartridges were then washed using 6 ml wash solution and peptides were eluted in 3 ml activation solution. All samples were dried overnight using a Genevac**™** MiVac concentrator at 37°C and stored at 4°C until use. Samples were resuspended in 200 µl ddH_2_O immediately before use. Controls included phosphate-buffered saline (PBS; 150 mM NaCl, 0.025 M NaH_2_PO_4_.2H_2_O, 0.075 M Na_2_HPO_4,_ pH 7.4) spiked with synthetic As-FLP-18A (PBS-As-FLP-18 SB; 100 µM) before C18 perfusion, PBS spiked with synthetic As-FLP-18A after C18 perfusion (PBS-As-FLP-18 SA; 100 µM), and perfused activation solution only (resuspended in ddH_2_O; negative control). 4 or 40 µl As-PCF [15X concentrated; final concentration in recording chamber 0.015X and 0.15X As-PCF respectively (where 1X As-PCF represents the native biological sample)], PBS-As-FLP-18 SB (final concentration in recording chamber 0.1 µM or 1 µM), or PBS-As-FLP-18 SA (final concentration in recording chamber 0.1 µM or 1 µM) were added to the recording chamber (n=4) and effects on ovijector muscle recorded. All data were analyzed as described [41]. Statistical analyses included repeated measures ANOVA and Student’s *t*-test to assess the significance of test substance/peptide effects and effect reversal after washout, respectively (see Fig 2).

### Cell-culture activation assays for *Caenorhabditis elegans* receptors

GPCR activation assay was performed as previously described [17, 53]. Briefly, mammalian CHO K1 cells stably overexpressing apo-aequorin and human Gα_16_ (ES-000-A24, PerkinElmer) were transiently transfected with *Ce-npr-*4a/pcDNA3.1, *Ce-npr-*5a/pcDNA3.1 or empty pcDNA3.1 (negative control) plasmid using Lipofectamine LTX and Plus reagent (Invitrogen, UK). After transfection, cells were grown in culture flasks overnight at 37 °C, after which they were transferred to 28 °C and allowed to incubate for 24 h. On the day of the assay, CHO cells were harvested and loaded with Coelenterazine H (Invitrogen) for 4 h at room temperature, which elicits the formation of the Ca^2+^-sensitive photoprotein aequorin. As-PCF or synthetic peptides (As-FLP-18 or Ce-FLP-18) dissolved in DMEM/BSA medium were added to cells and luminescence measured for 30 s at 496 nm using a Mithras LB940 luminometer (Berthold Technologies, Germany). After 30 s of readout, 0.1% Triton X-100 was added to lyse the cells, resulting in a maximal Ca^2+^ response that was measured for 10 s. Note that *C. elegans* synthetic peptides were synthesized by Thermo Scientific and *A. suum* synthetic peptides were synthesized by Genosphere Biotechnologies (France), all peptides were tested at a final concentration of 10 μM. The final concentration of As-PCF was half the concentration of the undiluted sample. In addition, a 10-fold dilution series ranging from 1/20 to 1/2000 of the undiluted As-PCF sample was tested. Assays were performed in triplicate on at least two different days.

### *Caenorhabditis elegans* bioassays

#### Body length assay

*C. elegans* body size assays were carried out as previously described [45]. Nematodes were treated with either 1, 2 or 5% As-PCF [diluted in S-medium, 1 L S Basal, 10 ml 1 M potassium citrate pH 6.0, 10 ml trace metals solution, 3 ml 1 M CaCl_2_, 3 ml 1 M MgSO_4_) [52]],10 µM synthetic As-FLP-18A [positive control; Genosphere Biotechnologies (France)] or S-medium (negative control; n>9 nematodes per treatment). Following incubation at 20°C for 96 hr nematodes were immobilized in M9 buffer containing 50 mM sodium azide (NaN_3_) and body length was calculated using a Leica MZ 12.5 stereomicroscope, Unibrain Fire-i digital camera and ImageJ software [54]. *Brood size assay:* Five L4 *C. elegans* were transferred into one well of a 24-well plate containing either 1, 2 or 5% As-PCF (diluted in S-medium), 10 µM synthetic As-FLP-18A (positive control) or S-medium (negative control) and supplemented with *E. coli* OP50 to a final volume of 1 ml (n=4 wells/treatment). Brood size was calculated by counting the number of progeny in each well after 96 hr at 21°C [46]. *wMicroTracker motility assay:* 50 L4 *C. elegans* in M9 buffer were added to each well of a 96-well plate along with *E. coli* OP50 (OD_600nm_ of 1) to a final volume of 90 µl. Basal locomotory activity was recorded for 1 hr using the wMicroTracker (InVivo Biosystems, Oregon, USA) with a bin size of 30 mins. After 1 hr each well was supplemented with 10 µl 1, 2, 5, 7.5 or 10% As-PCF (diluted in M9), M9 buffer (negative control) or 10 uM synthetic As-FLP-18A (positive control) (n>4 wells/treatment). Nematode locomotory activity was recorded for a further 12 hrs. Data were analyzed using GraphPad Prism version 8. Statistical significance was determined by one-way ANOVA and Dunnett’s multiple comparisons test or two-way ANOVA and Tukey’s multiple comparisons test.

### *In silico* identification of *flp-18, npr-4* and *-5*

Putative nematode *flp-18, npr-4* and *npr-5* orthologs were identified from WormBase ParaSite version 14 [https://parasite.wormbase.org/index.html; 49, 50, 51] and aligned using CLUSTAL Omega [default settings; 55]. Alignments were subsequently used to construct Hidden Markov Models (HMM) using HMMER v3.2 [default hmmbuild parameters; 56] based on methods previously described [57]. *hmmsearch* (default settings) was employed to identify potential *flp-18, npr-4* and *npr-5* sequences within the predicted protein datasets of 109 nematode species [134 genomes, see Supplementary Data 2 (SD2); 51]. The putative *npr-4* and *-5* sequences identified via *hmmsearch* were then used as queries in BLASTp searches in the NCBI non-redundant *C. elegans* predicted protein database (https://blast.ncbi.nlm.nih.gov; default settings). The putative *flp-18* sequences identified via *hmmsearch* were confirmed visually via the presence of the conserved PGXXRFG C-terminal motif. Queries that failed to return a putative target gene as the highest scoring pair/top hit were excluded from downstream analyses. Genomes lacking putative target gene hits were subjected to manual BLASTp and tBLASTn using WormBase ParaSite version 14 and *C. elegans flp-18, npr-4* and *npr-5* as query sequences. Additionally, orthologs from the most closely related nematode species were also used as queries in BLASTp/ tBLASTn searches.

### Statistical analysis

All statistical analyses were performed using GraphPad Prism Version 8. Statistical tests included one-way ANOVA (Dunnett’s multiple comparisons test), two-way ANOVA (Tukey’s multiple comparisons test), repeated measures ANOVA and Student’s *t*-test. LC-MS/MS data were analyzed using PEAKS Studio 7 (Bioinformatics Solution Inc., Waterloo, ON, Canada)

## Supporting information

Supplemental References

## Acknowledgments

Authors thank Dr Brett Greer for technical assistance/equipment and Sorcha Donnelly for assistance with As-PCF collection. The authors are grateful to Karro, Cookstown, NI for the assistance in the collection of nematodes. The authors acknowledge support for this work from: Biotechnology and Biological Sciences Research Council/Boehringer Ingelheim grant BB/MO10392/1 (AM, NJM and AGM); Department for Education Northern Ireland (DfE) Studentship (FMcK); Department of Agriculture Environment and Rural Affairs Northern Ireland (DAERA) Studentship (AI); National Institutes of Health (NIH, USA) grant GM097435 (MM); National Institutes of Health (NIH, USA) grants R01DK071801 and S10RR029531 (LL); Research Foundation Flanders (FWO) grant G0C0618N (IB).

## Supplementary information

**S1 File. As-PCF peptides detected in this study**. High confidence peptides (>1% FDR) detected across all samples (Table 1; Tab 1); All peptides detected in single female samples (Table 2; Tab 2); All peptides detected in pooled female samples (Table 3; Tab 3); All peptides detected in pooled male samples (Table 4; Tab 4). In all cases −10lgP denotes P-value [-10*log10(P-value)] as converted by PEAKS software.

**S2 File. Nematode genomes mined for gene sequelogues of *flp*-18, *npr*-4 and *npr*-5**. Wormbase ParaSite [https://parasite.wormbase.org/index.html; 49, 50, 51] gene IDs are provided for each identified sequelogue. Gene IDs highlighted in red indicate sequences that were used to build Hidden Markov Models. Genomic locations for representative partial sequences from TBLASTN hits are detailed where appropriate.

**S1 Table. The modulatory effects of As-PCF on *Ascaris suum* ovijector tissue preparations**.

**S2 Table. The modulatory effects of synthetic As-FLP-18A on *Ascaris suum* ovijector tissue preparations**.

## References

1. White JG, Southgate E, Thomson JN, Brenner S. The structure of the nervous system of the nematode Caenorhabditis elegans. Philos Trans R Soc Lond B Biol Sci. 1986;314(1165):1–340. Epub 1986/11/12. doi: 10.1098/rstb.1986.0056. PubMed PMID: 22462104.

2. Cook SJ, Jarrell TA, Brittin CA, Wang Y, Bloniarz AE, Yakovlev MA, et al. Whole-animal connectomes of both Caenorhabditis elegans sexes. Nature. 2019;571(7763):63–71. Epub 2019/07/05. doi: 10.1038/s41586-019-1352-7. PubMed PMID: 31270481.

3. Albertson DG, Thomson JN. The pharynx of Caenorhabditis elegans. Philos Trans R Soc Lond B Biol Sci. 1976;275(938):299–325. Epub 1976/08/10. doi: 10.1098/rstb.1976.0085. PubMed PMID: 8805.

4. Stretton AOW, Maule AG. Chapter 6 – The Neurobiology of Ascaris and Other Parasitic Nematodes. In: Holland C, editor. Ascaris: The Neglected Parasite. Amsterdam: Elsevier; 2013. p. 127–52.

5. Bargmann CI. Beyond the connectome: how neuromodulators shape neural circuits. Bioessays. 2012;34(6):458–65. Epub 2012/03/08. doi: 10.1002/bies.201100185. PubMed PMID: 22396302.

6. Brezina V. Beyond the wiring diagram: signalling through complex neuromodulator networks. Philosophical transactions of the Royal Society of London Series B, Biological sciences. 2010;365(1551):2363–74. doi: 10.1098/rstb.2010.0105. PubMed PMID: 20603357.

7. Liang Z, Schmerberg CM, Li L. Mass spectrometric measurement of neuropeptide secretion in the crab, Cancer borealis, by in vivo microdialysis. Analyst. 2015;140(11):3803–13. Epub 2014/12/30. doi: 10.1039/c4an02016b. PubMed PMID: 25537886; PubMed Central PMCID: PMCPMC4837892.

8. Jekely G, Melzer S, Beets I, Kadow ICG, Koene J, Haddad S, et al. The long and the short of it – a perspective on peptidergic regulation of circuits and behaviour. J Exp Biol. 2018;221(Pt 3). Epub 2018/02/14. doi: 10.1242/jeb.166710. PubMed PMID: 29439060.

9. Smart D, Shaw C, Johnston CF, Halton DW, Fairweather I, Buchanan KD. Chromatographic and immunological characterisation of neuropeptide Y-like and pancreatic polypeptide-like peptides from the nematode Ascaris suum. Comp Biochem Physiol C. 1992;102(3):477–81. Epub 1992/07/01. PubMed PMID: 1360356.

10. Laurent P, Ch’ng Q, Jospin M, Chen C, Lorenzo R, de Bono M. Genetic dissection of neuropeptide cell biology at high and low activity in a defined sensory neuron. Proc Natl Acad Sci U S A. 2018;115(29):E6890–e9. Epub 2018/07/01. doi: 10.1073/pnas.1714610115. PubMed PMID: 29959203; PubMed Central PMCID: PMCPMC6055185.

11. Wang H, Girskis K, Janssen T, Chan JP, Dasgupta K, Knowles JA, et al. Neuropeptide secreted from a pacemaker activates neurons to control a rhythmic behavior. Curr Biol. 2013;23(9):746–54. Epub 2013/04/16. doi: 10.1016/j.cub.2013.03.049. PubMed PMID: 23583549; PubMed Central PMCID: PMCPMC3651789.

12. Sieburth D, Madison JM, Kaplan JM. PKC-1 regulates secretion of neuropeptides. Nat Neurosci. 2007;10(1):49–57. Epub 2006/11/28. doi: 10.1038/nn1810. PubMed PMID: 17128266.

13. Rogers C, Reale V, Kim K, Chatwin H, Li C, Evans P, et al. Inhibition of Caenorhabditis elegans social feeding by FMRFamide-related peptide activation of NPR-1. Nat Neurosci. 2003;6(11):1178–85. Epub 2003/10/14. doi: 10.1038/nn1140. PubMed PMID: 14555955.

14. Komuniecki R, Hapiak V, Harris G, Bamber B. Context-dependent modulation reconfigures interactive sensory-mediated microcircuits in Caenorhabditis elegans. Curr Opin Neurobiol. 2014;29:17–24. Epub 2014/05/09. doi: 10.1016/j.conb.2014.04.006. PubMed PMID: 24811318.

15. Chase DL, Koelle MR. Biogenic amine neurotransmitters in C. elegans. WormBook. 2007:1–15. Epub 2007/12/01. doi: 10.1895/wormbook.1.132.1. PubMed PMID: 18050501; PubMed Central PMCID: PMCPMC4781333.

16. Chew YL, Tanizawa Y, Cho Y, Zhao B, Yu AJ, Ardiel EL, et al. An Afferent Neuropeptide System Transmits Mechanosensory Signals Triggering Sensitization and Arousal in C. elegans. Neuron. 2018;99(6):1233–46.e6. Epub 2018/08/28. doi: 10.1016/j.neuron.2018.08.003. PubMed PMID: 30146306; PubMed Central PMCID: PMCPMC6162336.

17. Beets I, Janssen T, Meelkop E, Temmerman L, Suetens N, Rademakers S, et al. Vasopressin/oxytocin-related signaling regulates gustatory associative learning in C. elegans. Science. 2012;338(6106):543–5. Epub 2012/11/01. doi: 10.1126/science.1226860. PubMed PMID: 23112336.

18. Chase DL, Pepper JS, Koelle MR. Mechanism of extrasynaptic dopamine signaling in Caenorhabditis elegans. Nat Neurosci. 2004;7(10):1096–103. Epub 2004/09/21. doi: 10.1038/nn1316. PubMed PMID: 15378064.

19. Bentley B, Branicky R, Barnes CL, Chew YL, Yemini E, Bullmore ET, et al. The Multilayer Connectome of Caenorhabditis elegans. PLoS Comput Biol. 2016;12(12):e1005283. Epub 2016/12/17. doi: 10.1371/journal.pcbi.1005283. PubMed PMID: 27984591; PubMed Central PMCID: PMCPMC5215746 GlaxoSmithKline; he holds stock in GSK. The authors have declared that no competing interests exist.

20. McCoy CJ, Atkinson LE, Robb E, Marks NJ, Maule AG, Mousley A. Tool-Driven Advances in Neuropeptide Research from a Nematode Parasite Perspective. Trends Parasitol. 2017;33(12):986–1002. Epub 2017/10/08. doi: 10.1016/j.pt.2017.08.009. PubMed PMID: 28986106.

21. Zhang J, Xin L, Shan B, Chen W, Xie M, Yuen D, et al. PEAKS DB: de novo sequencing assisted database search for sensitive and accurate peptide identification. Mol Cell Proteomics. 2012;11(4):M111.010587. Epub 2011/12/22. doi: 10.1074/mcp.M111.010587. PubMed PMID: 22186715; PubMed Central PMCID: PMCPMC3322562.

22. Jex AR, Liu S, Li B, Young ND, Hall RS, Li Y, et al. Ascaris suum draft genome. Nature. 2011;479(7374):529–33. Epub 2011/10/28. doi: 10.1038/nature10553. PubMed PMID: 22031327.

23. Pillai A, Ueno S, Zhang H, Lee JM, Kato Y. Cecropin P1 and novel nematode cecropins: a bacteria-inducible antimicrobial peptide family in the nematode Ascaris suum. Biochem J. 2005;390(Pt 1):207–14. Epub 2005/04/27. doi: 10.1042/BJ20050218. PubMed PMID: 15850460; PubMed Central PMCID: PMCPMC1184576.

24. Nusbaum MP, Blitz DM, Marder E. Functional consequences of neuropeptide and small-molecule co-transmission. Nat Rev Neurosci. 2017;18(7):389–403. Epub 2017/06/09. doi: 10.1038/nrn.2017.56. PubMed PMID: 28592905; PubMed Central PMCID: PMCPMC5547741.

25. Ch’ng Q, Sieburth D, Kaplan JM. Profiling synaptic proteins identifies regulators of insulin secretion and lifespan. PLoS Genet. 2008;4(11):e1000283. Epub 2008/12/02. doi: 10.1371/journal.pgen.1000283. PubMed PMID: 19043554; PubMed Central PMCID: PMCPMC2582949.

26. Hao Y, Hu Z, Sieburth D, Kaplan JM. RIC-7 promotes neuropeptide secretion. PLoS Genet. 2012;8(1):e1002464. Epub 2012/01/26. doi: 10.1371/journal.pgen.1002464. PubMed PMID: 22275875; PubMed Central PMCID: PMCPMC3261915.

27. Persson MG, Eklund MB, Dircksen H, Muren JE, Nassel DR. Pigment-dispersing factor in the locust abdominal ganglia may have roles as circulating neurohormone and central neuromodulator. J Neurobiol. 2001;48(1):19–41. Epub 2001/06/08. PubMed PMID: 11391647.

28. Choi S, Chatzigeorgiou M, Taylor KP, Schafer WR, Kaplan JM. Analysis of NPR-1 reveals a circuit mechanism for behavioral quiescence in C. elegans. Neuron. 2013;78(5):869–80. Epub 2013/06/15. doi: 10.1016/j.neuron.2013.04.002. PubMed PMID: 23764289; PubMed Central PMCID: PMCPMC3683153.

29. Janssen T, Husson SJ, Lindemans M, Mertens I, Rademakers S, Ver Donck K, et al. Functional characterization of three G protein-coupled receptors for pigment dispersing factors in Caenorhabditis elegans. J Biol Chem. 2008;283(22):15241–9. Epub 2008/04/09. doi: 10.1074/jbc.M709060200. PubMed PMID: 18390545; PubMed Central PMCID: PMCPMC3258896.

30. Janssen T, Husson SJ, Meelkop E, Temmerman L, Lindemans M, Verstraelen K, et al. Discovery and characterization of a conserved pigment dispersing factor-like neuropeptide pathway in Caenorhabditis elegans. J Neurochem. 2009;111(1):228–41. Epub 2009/08/19. doi: 10.1111/j.1471-4159.2009.06323.x. PubMed PMID: 19686386.

31. Cohen M, Reale V, Olofsson B, Knights A, Evans P, de Bono M. Coordinated regulation of foraging and metabolism in C. elegans by RFamide neuropeptide signaling. Cell Metab. 2009;9(4):375–85. Epub 2009/04/10. doi: 10.1016/j.cmet.2009.02.003. PubMed PMID: 19356718.

32. Rhoads ML, Fetterer RH. Purification and characterisation of a secreted aminopeptidase from adult Ascaris suum. Int J Parasitol. 1998;28(11):1681–90. Epub 1998/12/10. doi: 10.1016/s0020-7519(98)00091-5. PubMed PMID: 9846604.

33. Wang T, Van Steendam K, Dhaenens M, Vlaminck J, Deforce D, Jex AR, et al. Proteomic analysis of the excretory-secretory products from larval stages of Ascaris suum reveals high abundance of glycosyl hydrolases. PLoS Negl Trop Dis. 2013;7(10):e2467. Epub 2013/10/08. doi: 10.1371/journal.pntd.0002467. PubMed PMID: 24098821; PubMed Central PMCID: PMCPMC3789772.

34. Chehayeb JF, Robertson AP, Martin RJ, Geary TG. Proteomic analysis of adult Ascaris suum fluid compartments and secretory products. PLoS Negl Trop Dis. 2014;8(6):e2939. Epub 2014/06/06. doi: 10.1371/journal.pntd.0002939. PubMed PMID: 24901219; PubMed Central PMCID: PMCPMC4046973.

35. Rosa BA, Townsend R, Jasmer DP, Mitreva M. Functional and phylogenetic characterization of proteins detected in various nematode intestinal compartments. Mol Cell Proteomics. 2015;14(4):812–27. Epub 2015/01/23. doi: 10.1074/mcp.M114.046227. PubMed PMID: 25609831; PubMed Central PMCID: PMCPMC4390262.

36. Nassel DR. Neuropeptide signaling near and far: how localized and timed is the action of neuropeptides in brain circuits? Invert Neurosci. 2009;9(2):57–75. Epub 2009/09/17. doi: 10.1007/s10158-009-0090-1. PubMed PMID: 19756790.

37. Fellowes RA, Dougan PM, Maule AG, Marks NJ, Halton DW. Neuromusculature of the ovijector of ascaris suum (Ascaroidea, nematoda): an ultrastructural and immunocytochemical study. J Comp Neurol. 1999;415(4):518–28. Epub 1999/11/26. doi: 10.1002/(sici)1096-9861(19991227)415:4<518::aid-cne7>3.0.co;2-l. PubMed PMID: 10570459.

38. Moffett CL, Beckett AM, Mousley A, Geary TG, Marks NJ, Halton DW, et al. The ovijector of Ascaris suum: multiple response types revealed by Caenorhabditis elegans FMRFamide-related peptides. International Journal for Parasitology. 2003;33(8):859–76. doi: 10.1016/s0020-7519(03)00109-7.

39. McVeigh P, Geary TG, Marks NJ, Maule AG. The FLP-side of nematodes. Trends Parasitol. 2006;22(8):385–96. Epub 2006/07/11. doi: 10.1016/j.pt.2006.06.010. PubMed PMID: 16824799.

40. Atkinson LE, Miskelly IR, Moffett CL, McCoy CJ, Maule AG, Marks NJ, et al. Unraveling flp-11/flp-32 dichotomy in nematodes. Int J Parasitol. 2016;46(11):723–36. Epub 2016/07/28. doi: 10.1016/j.ijpara.2016.05.010. PubMed PMID: 27451358; PubMed Central PMCID: PMCPMC5038847.

41. Fellowes RA, Maule AG, Marks NJ, Geary TG, Thompson DP, Shaw C, et al. Modulation of the motility of the vagina vera of Ascaris suum in vitro by FMRF amide-related peptides. Parasitology. 1998;116 (Pt 3):277–87. Epub 1998/04/29. doi: 10.1017/s0031182097002229. PubMed PMID: 9550221.

42. Fellowes RA, Maule AG, Martin RJ, Geary TG, Thompson DP, Kimber MJ, et al. Classical neurotransmitters in the ovijector of Ascaris suum: localization and modulation of muscle activity. Parasitology. 2000;121 (Pt 3):325–36. Epub 2000/11/21. doi: 10.1017/s0031182099006290. PubMed PMID: 11085252.

43. Nanda JC, Stretton AO. In situ hybridization of neuropeptide-encoding transcripts afp-1, afp-3, and afp-4 in neurons of the nematode Ascaris suum. J Comp Neurol. 2010;518(6):896–910. Epub 2010/01/09. doi: 10.1002/cne.22251. PubMed PMID: 20058230; PubMed Central PMCID: PMCPMC2972677.

44. Anderson RC, Newton CL, Millar RP, Katz AA. The Brugia malayi neuropeptide receptor-4 is activated by FMRFamide-like peptides and signals via Gαi. Mol Biochem Parasitol. 2014;195(1):54–8. Epub 2014/07/20. doi: 10.1016/j.molbiopara.2014.07.002. PubMed PMID: 25038481.

45. Nagashima T, Oami E, Kutsuna N, Ishiura S, Suo S. Dopamine regulates body size in Caenorhabditis elegans. Dev Biol. 2016;412(1):128–38. Epub 2016/02/28. doi: 10.1016/j.ydbio.2016.02.021. PubMed PMID: 26921458.

46. Bansal A, Zhu LJ, Yen K, Tissenbaum HA. Uncoupling lifespan and healthspan in Caenorhabditis elegans longevity mutants. Proc Natl Acad Sci U S A. 2015;112(3):E277–86. Epub 2015/01/07. doi: 10.1073/pnas.1412192112. PubMed PMID: 25561524; PubMed Central PMCID: PMCPMC4311797.

47. Kage-Nakadai E, Kobuna H, Kimura M, Gengyo-Ando K, Inoue T, Arai H, et al. Two very long chain fatty acid acyl-CoA synthetase genes, acs-20 and acs-22, have roles in the cuticle surface barrier in Caenorhabditis elegans. PLoS One. 2010;5(1):e8857. Epub 2010/01/30. doi: 10.1371/journal.pone.0008857. PubMed PMID: 20111596; PubMed Central PMCID: PMCPMC2810326.

48. Li C, Kim K. Family of FLP Peptides in Caenorhabditis elegans and Related Nematodes. Frontiers in Endocrinology. 2014;5(150). doi: 10.3389/fendo.2014.00150.

49. Howe KL, Bolt BJ, Cain S, Chan J, Chen WJ, Davis P, et al. WormBase 2016: expanding to enable helminth genomic research. Nucleic Acids Res. 2016;44(D1):D774–80. Epub 2015/11/19. doi: 10.1093/nar/gkv1217. PubMed PMID: 26578572; PubMed Central PMCID: PMCPMC4702863.

50. Bolt BJ, Rodgers FH, Shafie M, Kersey PJ, Berriman M, Howe KL. Using WormBase ParaSite: An Integrated Platform for Exploring Helminth Genomic Data. Methods Mol Biol. 2018;1757:471–91. Epub 2018/05/16. doi: 10.1007/978-1-4939-7737-6_15. PubMed PMID: 29761467.

51. Howe KL, Bolt BJ, Shafie M, Kersey P, Berriman M. WormBase ParaSite – a comprehensive resource for helminth genomics. Mol Biochem Parasitol. 2017;215:2–10. Epub 2016/12/03. doi: 10.1016/j.molbiopara.2016.11.005. PubMed PMID: 27899279; PubMed Central PMCID: PMCPMC5486357.

52. Stiernagle T. Maintenance of C. elegans. WormBook. 2006:1–11. Epub 2007/12/01. doi: 10.1895/wormbook.1.101.1. PubMed PMID: 18050451; PubMed Central PMCID: PMCPMC4781397.

53. Van Sinay E, Mirabeau O, Depuydt G, Van Hiel MB, Peymen K, Watteyne J, et al. Evolutionarily conserved TRH neuropeptide pathway regulates growth in Caenorhabditis elegans. Proc Natl Acad Sci U S A. 2017;114(20):E4065–e74. Epub 2017/05/04. doi: 10.1073/pnas.1617392114. PubMed PMID: 28461507; PubMed Central PMCID: PMCPMC5441806.

54. Schneider CA, Rasband WS, Eliceiri KW. NIH Image to ImageJ: 25 years of image analysis. Nat Methods. 2012;9(7):671–5. Epub 2012/08/30. doi: 10.1038/nmeth.2089. PubMed PMID: 22930834; PubMed Central PMCID: PMCPMC5554542.

55. Chojnacki S, Cowley A, Lee J, Foix A, Lopez R. Programmatic access to bioinformatics tools from EMBL-EBI update: 2017. Nucleic Acids Res. 2017;45(W1):W550–w3. Epub 2017/04/22. doi: 10.1093/nar/gkx273. PubMed PMID: 28431173; PubMed Central PMCID: PMCPMC5570243.

56. Mistry J, Finn RD, Eddy SR, Bateman A, Punta M. Challenges in homology search: HMMER3 and convergent evolution of coiled-coil regions. Nucleic Acids Res. 2013;41(12):e121. Epub 2013/04/20. doi: 10.1093/nar/gkt263. PubMed PMID: 23598997; PubMed Central PMCID: PMCPMC3695513.

57. McVeigh P, McCammick E, McCusker P, Wells D, Hodgkinson J, Paterson S, et al. Profiling G protein-coupled receptors of Fasciola hepatica identifies orphan rhodopsins unique to phylum Platyhelminthes. Int J Parasitol Drugs Drug Resist. 2018;8(1):87–103. Epub 2018/02/24. doi: 10.1016/j.ijpddr.2018.01.001. PubMed PMID: 29474932; PubMed Central PMCID: PMCPMC6114109.

58. Holterman M, van der Wurff A, van den Elsen S, van Megen H, Bongers T, Holovachov O, et al. Phylum-wide analysis of SSU rDNA reveals deep phylogenetic relationships among nematodes and accelerated evolution toward crown Clades. Mol Biol Evol. 2006;23(9):1792–800. Epub 2006/06/23. doi: 10.1093/molbev/msl044. PubMed PMID: 16790472.

59. Simao FA, Waterhouse RM, Ioannidis P, Kriventseva EV, Zdobnov EM. BUSCO: assessing genome assembly and annotation completeness with single-copy orthologs. Bioinformatics. 2015;31(19):3210–2. Epub 2015/06/11. doi: 10.1093/bioinformatics/btv351. PubMed PMID: 26059717.

60. Parra G, Bradnam K, Korf I. CEGMA: a pipeline to accurately annotate core genes in eukaryotic genomes. Bioinformatics. 2007;23(9):1061–7. Epub 2007/03/03. doi: 10.1093/bioinformatics/btm071. PubMed PMID: 17332020.

