## Supplemental Table 1 for "Extrasynaptic volume transmission: A novel route for neuropeptide signaling in nematodes"

**S1 Table. The modulatory effects of As-PCF on *Ascaris suum* ovijector tissue preparations.**

| **Test** | **Time**  (p.a.) | **Contraction freq.**  (% 0 min, 2 min  prior to min p.a.) | **Contraction ampl.**  (% 0 min, 2 min  prior to min p.a.) | **Change in tension**  (mg/mm, 5 min p.a.) |
| --- | --- | --- | --- | --- |
| **4 µl As-PCF**  n=7 | 10 secs  20 secs  30 secs  50 secs  70 secs  2 min  5 min  10 min  **OVERALL** | N/A  N/A  N/A  N/A  N/A  ↓ 58.16±12.33 🗸  ↓ 10.57±7.699 🗸  ↓ 3.810±3.810 🗸  **P<0.0001, F=43.61** | N/A  N/A  N/A  N/A  N/A  ↓ 96.25±7.177 🗴  ↓ 23.85±17.36 🗸  ↓ 16.21±16.21 🗸  **P=0.0151, F=3.647** | Sht 0.002±0.01  Sht 0.001±0.01  Lng -0.02±0.01  Lng -0.02±0.01  Lng -0.001±0.02  Sht 0.04±0.05  Sht 0.08±0.05  Sht 0.11±0.06  **NS** |
| **40 µl As-PCF**  n=7 | 10 secs  20 secs  30 secs  50 secs  70 secs  2 min  5 min  10 min  **OVERALL** | N/A  N/A  N/A  N/A  N/A  ↓ 46.48±9.588 🗸  ↓ Ceased 🗸  ↓ Ceased 🗸  **P<0.0001, F=98.56** | N/A  N/A  N/A  N/A  N/A  ↑ 131.1±23.72 🗴  ↓ Ceased 🗸  ↓ Ceased 🗸  **P<0.0001, F=32.80** | Lng -0.02±0.04  Lng -0.05±0.05  Lng -0.08±0.08  Lng -0.08±0.10  Sht 0.05±0.09  Sht 0.10±0.07  Sht 0.15±0.05  Sht 0.20±0.06  **NS** |
| **4 µl SB**  **(As-FLP-18A)**  n=6 | 10 secs  20 secs  30 secs  50 secs  70 secs  2 min  5 min  10 min  **OVERALL** | N/A  N/A  N/A  N/A  N/A  ↓ 43.00±5.223 🗸  ↓ 8.730±7.817 🗸  ↓ Ceased 🗸  **P<0.0001, F=91.35** | N/A  N/A  N/A  N/A  N/A  ↑ 154.5±19.98  ↓ 81.52±70.6  ↓ Ceased  **NS** | 0.00±0.00  Lng -0.01±0.01  Lng -0.06±0.04  Lng -0.13±0.07  Lng -0.11±0.07  Lng -0.04±0.06  Sht 0.03±0.04  Sht 0.05±0.04  **NS** |
| **40 µl SB**  **(As-FLP-18A)**  n=3 | 10 secs  20 secs  30 secs  50 secs  70 secs  2 min  5 min  10 min  **OVERALL** | N/A  N/A  N/A  N/A  N/A  ↓ 6.754±0.8874 🗸  ↓ Ceased 🗸  ↓ Ceased 🗸  **P<0.0001, F=12186** | N/A  N/A  N/A  N/A  N/A  ↑ 389.2±171.3  ↓ Ceased  ↓ Ceased  **NS** | Lng -0.26±0.08  Lng -0.26±0.08  Lng -0.23±0.09  Lng -0.22±0.09  Lng -0.20±0.10  Lng -0.18±0.12  Lng -0.10±0.10  Lng -0.04±0.07  **NS** |
| **4 µl SA**  **(As-FLP-18A)**  n=5 | 10 secs  20 secs  30 secs  50 secs  70 secs  2 min  5 min  10 min  **OVERALL** | N/A  N/A  N/A  N/A  N/A  ↓ 84.28±15.31 🗴  ↓ 52.98±25.80 🗴  ↓ 34.48±21.11 🗸  **P=0.0149, F=5.285** | N/A  N/A  N/A  N/A  N/A  ↑ 119.3±15.54 🗴  ↓ 68.72±19.68 🗴  ↓ 35.89±22.41 🗸  **P=0.0070, F=6.595** | Lng -0.01±0.01  Lng -0.02±0.01  Lng -0.03±0.02  Lng -0.04±0.04  Lng -0.002±0.05  Sht 0.008±0.06  Sht 0.04±0.06  Sht 0.05±0.04  **NS** |
| **40 µl SA**  **(As-FLP-18A)**  n=4 | 10 secs  20 secs  30 secs  50 secs  70 secs  2 min  5 min  10 min  **OVERALL** | N/A  N/A  N/A  N/A  N/A  ↓ 37.85±22.51 🗸  ↓ Ceased 🗸  ↓ 3.125±3.125 🗸  **P=0.0004, F=17.29** | N/A  N/A  N/A  N/A  N/A  ↓ 41.31±23.90 🗸  ↓ Ceased 🗸  ↓ 16.90±16.90 🗸  **P=0.0025, F=10.68** | Sht 0.03±0.06  Sht 0.02±0.07  Lng -0.001±0.09  Sht 0.05±0.07  Sht 0.07±0.05  Sht 0.09±0.04  Sht 0.11±0.02  Sht 0.10±0.02  **NS** |
| **Negative control**  (n=3) | 10 secs  20 secs  30 secs  50 secs  70 secs  2 min  5 min  10 min  **OVERALL** | N/A  N/A  N/A  N/A  N/A  ↑ 100.36±8.567  ↑ 112.02±5.802  ↑ 116.97±14.66  **NS** | N/A  N/A  N/A  N/A  N/A  ↑ 103.70±4.436  ↓ 92.63±5.689  ↓ 88.94±8.705  **NS** | N/A  N/A  N/A  N/A  N/A  0  0  Lng -0.011±0.01  **NS** |

% 0 min, values presented as percentage of that recorded at time 0; p.a., post-addition; ↓, denotes decrease; ↑, denotes increase; 🗴, denotes non-significant Dunnett’s post hoc test; 🗸, denotes significant Dunnett’s post hoc test; NS, not significant; Sht, shortening of tissue; Lng, lengthening of tissue**.**
