## Supplemental Table 2 for "Extrasynaptic volume transmission: A novel route for neuropeptide signaling in nematodes"

**S2 Table.** **The modulatory effects of synthetic As-FLP-18A on *Ascaris suum* ovijector tissue preparations.**

| **Test** | **Time**  (p.a.) | **Contraction freq.**  (% 0 min, 2 min  prior to min p.a.) | **Contraction ampl.**  (% 0 min, 2 min  prior to min p.a.) | **Change in tension**  (mg/mm, 5 min p.a.) |
| --- | --- | --- | --- | --- |
| **10 µM**  **As-FLP-18A**  n=13 | 10 secs  20 secs  30 secs  50 secs  70 secs  2 min  5 min  10 min  **OVERALL** | N/A  N/A  N/A  N/A  N/A  ↓ 52.32±9.931 🗸  ↓ 5.015±3.097 🗸  ↓ 4.872±3.897 🗸  **P<0.0001, F=67.76** | N/A  N/A  N/A  N/A  N/A  ↑ 111.4±11.55 🗴  ↓ 24.33±14.28 🗸  ↓ 11.51±8.354 🗸  **P<0.0001, F=25.54** | 0.00±0.00🗴  Sht _+_0.01±0.01 🗴  Lng -0.07±0.04 🗴  Sht +0.05±0.02 🗸  Sht +0.09±0.03 🗸  Sht +0.12±0.02 🗸  Sht +0.13±0.01 🗸  Lng -0.03±0.02 🗸  **P<0.0001, F=12.26** |
| **1 µM**  **As-FLP-18A**  n=6 | 10 secs  20 secs  30 secs  50 secs  70 secs  2 min  5 min  10 min  **OVERALL** | N/A  N/A  N/A  N/A  N/A  ↓ 83.01±16.35 🗴  ↓ 47.66±18.09 🗴  ↓ 34.08±16.39 🗸  **P=0.0489, F=3.521** | N/A  N/A  N/A  N/A  N/A  ↑ 126.2±18.34  ↑ 102.7±30.34  ↑ 119.0±34.18  **NS** | 0.00±0.00  0.00±0.00  Lng -0.11±0.07  Lng -0.13±0.08  Lng -0.14±0.08  Sht +0.02±0.06  Sht +0.03±0.03  Sht +0.002±0.02  **NS** |
| **0.1 µM**  **As-FLP-18A**  n=6 | 30 secs  2 min  5 min  10 min  **OVERALL** | N/A  ↓ 95.42±21.06 🗴  ↑128.8±11.92 🗴  ↑104.1± 18.30 🗴 **P=0.3250, F=1.282** | N/A  ↑ 130.3±12.89 🗴  ↑ 167.7±26.43 🗴  ↑ 224.5±56.79 🗸  **P=0.0340, F=4.025** | Lng -0.002±0.002 **🗴**  Lng -0.27±0.002 🗸  Lng -0.28±0.07 🗸  Lng -0.20±0.07 🗴  **P=0.0044, F=8.701** |
| 0.01 **µM**   **As-FLP-18A**  n=6 | 30 secs  2 min  5 min  10 min  **OVERALL** | N/A  ↑ 116.3±11.68 ↑ 109.6±11.67  ↓ 92.57±10.62  **NS** | N/A  ↑ 111.4±13.26  ↑ 152.8±41.98  ↑ 154.2±45.03  **NS** | Lng -0.04±0.03  Lng -0.07±0.03  Lng -0.08±0.04  Lng -0.05±0.05  **NS** |
| **1 nM**  **As-FLP-18A**  n=6 | 30 secs  2 min  5 min  10 min  **OVERALL** | N/A  ↑ 104.3±6.685  ↓ 90.13±6.395  ↓ 83.74±7.552  **NS** | N/A  ↓ 92.52±7.007  ↓ 97.22±5.835  ↑ 105.5±5.656  **NS** | Lng -0.01±0.01  Sht +0.01±0.01  Sht +0.01±0.01  Sht +0.03±0.01  **NS** |
| **-ve control**  (n=3) | 30 secs  2 min  5 min  10 min  **OVERALL** | N/A  ↓ 96.97±3.030  ↑ 101.96±1.961  ↑ 109.16±1.911  **NS** | N/A  ↓ 97.72±3.249  ↓ 91.04±2.207  ↓ 82.40±5.482  **NS** | N/A  0±0.00  0±0.00  0±0.00  **NS** |

% 0 min, values presented as percentage of that recorded at time 0; p.a., post-addition; ↓, denotes decrease; ↑, denotes increase; 🗴, denotes non-significant Dunnett’s post hoc test; 🗸, denotes significant Dunnett’s post hoc test; NS, not significant; Sht, shortening of tissue; Lng, lengthening of tissue.
