## Supplemental References for "Extrasynaptic volume transmission: A novel route for neuropeptide signaling in nematodes"

1.	Schiffer PH, Kroiher M, Kraus C, Koutsovoulos GD, Kumar S, Camps JIR, et al. The genome of Romanomermis culicivorax: revealing fundamental changes in the core developmental genetic toolkit in Nematoda. Bmc Genomics. 2013;14. doi: Unsp 923

10.1186/1471-2164-14-923. PubMed PMID: WOS:000329613800002.

2.	Coghlan A, Tyagi R, Cotton JA, Holroyd N, Rosa BA, Tsai IJ, et al. Comparative genomics of the major parasitic worms. Nat Genet. 2019;51(1):163-+. doi: 10.1038/s41588-018-0262-1. PubMed PMID: WOS:000454108800023.

3.	Korhonen PK, Pozio E, La Rosa G, Chang BCH, Koehler AV, Hoberg EP, et al. Phylogenomic and biogeographic reconstruction of the Trichinella complex. Nat Commun. 2016;7. doi: ARTN 10513

10.1038/ncomms10513. PubMed PMID: WOS:000371012700001.

4.	Mitreva M, Jasmer DP, Zarlenga DS, Wang Z, Abubucker S, Martin J, et al. The draft genome of the parasitic nematode Trichinella spiralis. Nat Genet. 2011;43(3):228-35. Epub 2011/02/22. doi: 10.1038/ng.769. PubMed PMID: 21336279; PubMed Central PMCID: PMCPMC3057868.

5.	Foth BJ, Tsai IJ, Reid AJ, Bancroft AJ, Nichol S, Tracey A, et al. Whipworm genome and dual-species transcriptome analyses provide molecular insights into an intimate host-parasite interaction. Nat Genet. 2014;46(7):693-700. doi: 10.1038/ng.3010. PubMed PMID: WOS:000338093800009.

6.	Harris TW, Arnaboldi V, Cain S, Chan J, Chen WJ, Cho J, et al. WormBase: a modern Model Organism Information Resource. Nucleic Acids Res. 2019. Epub 2019/10/24. doi: 10.1093/nar/gkz920. PubMed PMID: 31642470.

7.	Jex AR, Nejsum P, Schwarz EM, Hu L, Young ND, Hall RS, et al. Genome and transcriptome of the porcine whipworm Trichuris suis. Nat Genet. 2014;46(7):701-6. Epub 2014/06/16. doi: 10.1038/ng.3012. PubMed PMID: 24929829; PubMed Central PMCID: PMCPMC4105696.

8.	Rosic S, Amouroux R, Requena CE, Gomes A, Emperle M, Beltran T, et al. Evolutionary analysis indicates that DNA alkylation damage is a byproduct of cytosine DNA methyltransferase activity. Nat Genet. 2018;50(3). doi: 10.1038/s41588-018-0061-8. PubMed PMID: WOS:000427933400021.

9.	Wang JB, Mitreva M, Berriman M, Thorne A, Magrini V, Koutsovoulos G, et al. Silencing of Germline-Expressed Genes by DNA Elimination in Somatic Cells. Dev Cell. 2012;23(5):1072-80. doi: 10.1016/j.devcel.2012.09.020. PubMed PMID: WOS:000311134100021.

10.	Wang JB, Gao SH, Mostovoy Y, Kang YY, Zagoskin M, Sun YQ, et al. Comparative genome analysis of programmed DNA elimination in nematodes. Genome Research. 2017;27(12):2001-14. doi: 10.1101/gr.225730.117. PubMed PMID: WOS:000417047600004.

11.	Jex AR, Liu S, Li B, Young ND, Hall RS, Li Y, et al. Ascaris suum draft genome. Nature. 2011;479(7374):529-33. Epub 2011/10/28. doi: 10.1038/nature10553. PubMed PMID: 22031327.

12.	Ghedin E, Wang SL, Spiro D, Caler E, Zhao Q, Crabtree J, et al. Draft genome of the filarial nematode parasite Brugia malayi. Science. 2007;317(5845):1756-60. doi: 10.1126/science.1145406. PubMed PMID: WOS:000249585900050.

13.	Godel C, Kumar S, Koutsovoulos G, Ludin P, Nilsson D, Comandatore F, et al. The genome of the heartworm, Dirofilaria immitis, reveals drug and vaccine targets. Faseb J. 2012;26(11):4650-61. doi: 10.1096/fj.12-205096. PubMed PMID: WOS:000310574200026.

14.	Desjardins CA, Cerqueira GC, Goldberg JM, Hotopp JCD, Haas BJ, Zucker J, et al. Genomics of Loa loa, a Wolbachia-free filarial parasite of humans. Nat Genet. 2013;45(5):495-U55. doi: 10.1038/ng.2585. PubMed PMID: WOS:000318158200010.

15.	Tallon LJ, Liu XY, Bennuru S, Chibucos MC, Godinez A, Ott S, et al. Single molecule sequencing and genome assembly of a clinical specimen of Loa loa, the causative agent of loiasis. Bmc Genomics. 2014;15. doi: Artn 788

10.1186/1471-2164-15-788. PubMed PMID: WOS:000342096700002.

16.	Cotton JA, Bennuru S, Grote A, Harsha B, Tracey A, Beech R, et al. The genome of Onchocerca volvulus, agent of river blindness. Nat Microbiol. 2017;2(2). doi: UNSP 16216

10.1038/nmicrobiol.2016.216. PubMed PMID: WOS:000397104900014.

17.	Zhu XQ, Korhonen PK, Cai HM, Young ND, Nejsum P, von Samson-Himmelstjerna G, et al. Genetic blueprint of the zoonotic pathogen Toxocara canis. Nat Commun. 2015;6. doi: ARTN 6145

10.1038/ncomms7145. PubMed PMID: WOS:000350198500003.

18.	Small ST, Reimer LJ, Tisch DJ, King CL, Christensen BM, Siba PM, et al. Population genomics of the filarial nematode parasite Wuchereria bancrofti from mosquitoes. Mol Ecol. 2016;25(7):1465-77. doi: 10.1111/mec.13574. PubMed PMID: WOS:000373104500006.

19.	Schwarz EM, Hu Y, Antoshechkin I, Miller MM, Sternberg PW, Aroian RV. The genome and transcriptome of the zoonotic hookworm Ancylostoma ceylanicum identify infection-specific gene families. Nat Genet. 2015;47(4):416-22. Epub 2015/03/03. doi: 10.1038/ng.3237. PubMed PMID: 25730766; PubMed Central PMCID: PMCPMC4617383.

20.	Mortazavi A, Schwarz EM, Williams B, Schaeffer L, Antoshechkin I, Wold BJ, et al. Scaffolding a Caenorhabditis nematode genome with RNA-seq. Genome Res. 2010;20(12):1740-7. Epub 2010/10/29. doi: 10.1101/gr.111021.110. PubMed PMID: 20980554; PubMed Central PMCID: PMCPMC2990000.

21.	Stein LD, Bao Z, Blasiar D, Blumenthal T, Brent MR, Chen N, et al. The genome sequence of Caenorhabditis briggsae: a platform for comparative genomics. PLoS Biol. 2003;1(2):E45. Epub 2003/11/19. doi: 10.1371/journal.pbio.0000045. PubMed PMID: 14624247; PubMed Central PMCID: PMCPMC261899.

22.	Ross JA, Koboldt DC, Staisch JE, Chamberlin HM, Gupta BP, Miller RD, et al. Caenorhabditis briggsae recombinant inbred line genotypes reveal inter-strain incompatibility and the evolution of recombination. PLoS Genet. 2011;7(7):e1002174. Epub 2011/07/23. doi: 10.1371/journal.pgen.1002174. PubMed PMID: 21779179; PubMed Central PMCID: PMCPMC3136444.

23.	Consortium CeS. Genome sequence of the nematode C. elegans: a platform for investigating biology. Science. 1998;282(5396):2012-8. Epub 1998/12/16. doi: 10.1126/science.282.5396.2012. PubMed PMID: 9851916.

24.	Kanzaki N, Tsai IJ, Tanaka R, Hunt VL, Liu D, Tsuyama K, et al. Biology and genome of a newly discovered sibling species of Caenorhabditis elegans. Nat Commun. 2018;9(1):3216. Epub 2018/08/12. doi: 10.1038/s41467-018-05712-5. PubMed PMID: 30097582; PubMed Central PMCID: PMCPMC6086898.

25.	Fierst JL, Willis JH, Thomas CG, Wang W, Reynolds RM, Ahearne TE, et al. Reproductive Mode and the Evolution of Genome Size and Structure in Caenorhabditis Nematodes. Plos Genetics. 2015;11(6). doi: ARTN e1005323

10.1371/journal.pgen.1005323. PubMed PMID: WOS:000357341600048.

26.	Yin D, Schwarz EM, Thomas CG, Felde RL, Korf IF, Cutter AD, et al. Rapid genome shrinkage in a self-fertile nematode reveals sperm competition proteins. Science. 2018;359(6371):55-+. doi: 10.1126/science.aao0827. PubMed PMID: WOS:000419324700065.

27.	Stevens L, Felix MA, Beltran T, Braendle C, Caurcel C, Fausett S, et al. Comparative genomics of 10 new Caenorhabditis species. Evol Lett. 2019;3(2):217-36. Epub 2019/04/23. doi: 10.1002/evl3.110. PubMed PMID: 31007946; PubMed Central PMCID: PMCPMC6457397.

28.	Koutsovoulos G, Makepeace B, Tanya VN, Blaxter M. Palaeosymbiosis revealed by genomic fossils of Wolbachia in a strongyloidean nematode. PLoS Genet. 2014;10(6):e1004397. Epub 2014/06/06. doi: 10.1371/journal.pgen.1004397. PubMed PMID: 24901418; PubMed Central PMCID: PMCPMC4046930.

29.	McNulty SN, Strube C, Rosa BA, Martin JC, Tyagi R, Choi YJ, et al. Dictyocaulus viviparus genome, variome and transcriptome elucidate lungworm biology and support future intervention. Sci Rep. 2016;6:20316. Epub 2016/02/10. doi: 10.1038/srep20316. PubMed PMID: 26856411; PubMed Central PMCID: PMCPMC4746573.

30.	Hiraki H, Kagoshima H, Kraus C, Schiffer PH, Ueta Y, Kroiher M, et al. Genome analysis of Diploscapter coronatus: insights into molecular peculiarities of a nematode with parthenogenetic reproduction. BMC Genomics. 2017;18(1):478. Epub 2017/06/26. doi: 10.1186/s12864-017-3860-x. PubMed PMID: 28646875; PubMed Central PMCID: PMCPMC5483258.

31.	Fradin H, Kiontke K, Zegar C, Gutwein M, Lucas J, Kovtun M, et al. Genome Architecture and Evolution of a Unichromosomal Asexual Nematode. Curr Biol. 2017;27(19):2928-+. doi: 10.1016/j.cub.2017.08.038. PubMed PMID: WOS:000412561400022.

32.	Laing R, Kikuchi T, Martinelli A, Tsai IJ, Beech RN, Redman E, et al. The genome and transcriptome of Haemonchus contortus, a key model parasite for drug and vaccine discovery. Genome Biol. 2013;14(8):R88. Epub 2013/08/30. doi: 10.1186/gb-2013-14-8-r88. PubMed PMID: 23985316; PubMed Central PMCID: PMCPMC4054779.

33.	Schwarz EM, Korhonen PK, Campbell BE, Young ND, Jex AR, Jabbar A, et al. The genome and developmental transcriptome of the strongylid nematode Haemonchus contortus. Genome Biology. 2013;14(8). doi: ARTN R89

10.1186/gb-2013-14-8-r89. PubMed PMID: WOS:000328195400007.

34.	Bai X, Adams BJ, Ciche TA, Clifton S, Gaugler R, Kim KS, et al. A lover and a fighter: the genome sequence of an entomopathogenic nematode Heterorhabditis bacteriophora. Plos One. 2013;8(7):e69618. Epub 2013/07/23. doi: 10.1371/journal.pone.0069618. PubMed PMID: 23874975; PubMed Central PMCID: PMCPMC3715494.

35.	Grosmaire M, Launay C, Siegwald M, Brugiere T, Estrada-Virrueta L, Berger D, et al. Males as somatic investment in a parthenogenetic nematode. Science. 2019;363(6432):1210-3. Epub 2019/03/16. doi: 10.1126/science.aau0099. PubMed PMID: 30872523.

36.	Prabh N, Roeseler W, Witte H, Eberhardt G, Sommer RJ, Rodelsperger C. Deep taxon sampling reveals the evolutionary dynamics of novel gene families in Pristionchus nematodes. Genome Research. 2018;28(11):1664-74. doi: 10.1101/gr.234971.118. PubMed PMID: WOS:000448950400007.

37.	Tang YT, Gao X, Rosa BA, Abubucker S, Hallsworth-Pepin K, Martin J, et al. Genome of the human hookworm Necator americanus. Nat Genet. 2014;46(3):261-+. doi: 10.1038/ng.2875. PubMed PMID: WOS:000332036700010.

38.	Tyagi R, Joachim A, Ruttkowski B, Rosa BA, Martin JC, Hallsworth-Pepin K, et al. Cracking the nodule worm code advances knowledge of parasite biology and biotechnology to tackle major diseases of livestock. Biotechnol Adv. 2015;33(6 Pt 1):980-91. Epub 2015/06/01. doi: 10.1016/j.biotechadv.2015.05.004. PubMed PMID: 26026709; PubMed Central PMCID: PMCPMC4746232.

39.	Besnard F, Koutsovoulos G, Dieudonne S, Blaxter M, Felix MA. Toward Universal Forward Genetics: Using a Draft Genome Sequence of the Nematode Oscheius tipulae To Identify Mutations Affecting Vulva Development. Genetics. 2017;206(4):1747-61. doi: 10.1534/genetics.117.203521. PubMed PMID: WOS:000406952400004.

40.	Rodelsperger C, Neher RA, Weller AM, Eberhardt G, Witte H, Mayer WE, et al. Characterization of genetic diversity in the nematode Pristionchus pacificus from population-scale resequencing data. Genetics. 2014;196(4):1153-65. Epub 2014/01/21. doi: 10.1534/genetics.113.159855. PubMed PMID: 24443445; PubMed Central PMCID: PMCPMC3982705.

41.	Dieterich C, Clifton SW, Schuster LN, Chinwalla A, Delehaunty K, Dinkelacker I, et al. The Pristionchus pacificus genome provides a unique perspective on nematode lifestyle and parasitism. Nat Genet. 2008;40(10):1193-8. doi: 10.1038/ng.227. PubMed PMID: WOS:000259651000018.

42.	Rodelsperger C, Meyer JM, Prabh N, Lanz C, Bemm F, Sommer RJ. Single-Molecule Sequencing Reveals the Chromosome-Scale Genomic Architecture of the Nematode Model Organism Pristionchus pacificus. Cell Rep. 2017;21(3):834-44. Epub 2017/10/19. doi: 10.1016/j.celrep.2017.09.077. PubMed PMID: 29045848.

43.	Kikuchi T, Cotton JA, Dalzell JJ, Hasegawa K, Kanzaki N, McVeigh P, et al. Genomic insights into the origin of parasitism in the emerging plant pathogen Bursaphelenchus xylophilus. Plos Pathog. 2011;7(9):e1002219. Epub 2011/09/13. doi: 10.1371/journal.ppat.1002219. PubMed PMID: 21909270; PubMed Central PMCID: PMCPMC3164644.

44.	Weinstein DJ, Allen SE, Lau MCY, Erasmus M, Asalone KC, Walters-Conte K, et al. The genome of a subterrestrial nematode reveals adaptations to heat. Nat Commun. 2019;10(1):5268. Epub 2019/11/23. doi: 10.1038/s41467-019-13245-8. PubMed PMID: 31754114; PubMed Central PMCID: PMCPMC6872716.

45.	Srinivasan J, Dillman AR, Macchietto MG, Heikkinen L, Lakso M, Fracchia KM, et al. The Draft Genome and Transcriptome of Panagrellus redivivus Are Shaped by the Harsh Demands of a Free-Living Lifestyle. Genetics. 2013;193(4):1279-+. doi: 10.1534/genetics.112.148809. PubMed PMID: WOS:000316937300020.

46.	Hunt VL, Tsai IJ, Coghlan A, Reid AJ, Holroyd N, Foth BJ, et al. The genomic basis of parasitism in the Strongyloides clade of nematodes. Nat Genet. 2016;48(3):299-307. Epub 2016/02/02. doi: 10.1038/ng.3495. PubMed PMID: 26829753; PubMed Central PMCID: PMCPMC4948059.

47.	Serra L, Macchietto M, Macias-Munoz A, McGill CJ, Rodriguez IM, Rodriguez B, et al. Hybrid Assembly of the Genome of the Entomopathogenic Nematode Steinernema carpocapsae Identifies the X-Chromosome. G3-Genes Genom Genet. 2019;9(8):2687-97. doi: 10.1534/g3.119.400180. PubMed PMID: WOS:000479303300029.

48.	Dillman AR, Macchietto M, Porter CF, Rogers A, Williams B, Antoshechkin I, et al. Comparative genomics of Steinernema reveals deeply conserved gene regulatory networks. Genome Biology. 2015;16. doi: ARTN 200

10.1186/s13059-015-0746-6. PubMed PMID: WOS:000361452900001.

49.	Schiffer PH, Polsky AL, Cole AG, Camps JIR, Kroiher M, Silver DH, et al. The gene regulatory program of Acrobeloides nanus reveals conservation of phylum-specific expression. P Natl Acad Sci USA. 2018;115(17):4459-64. doi: 10.1073/pnas.1720817115. PubMed PMID: WOS:000430697500067.

50.	Zheng JS, Peng DH, Chen L, Liu HL, Chen F, Xu MC, et al. The Ditylenchus destructor genome provides new insights into the evolution of plant parasitic nematodes. P Roy Soc B-Biol Sci. 2016;283(1835). doi: ARTN 20160942

10.1098/rspb.2016.0942. PubMed PMID: WOS:000382430800010.

51.	Mimee B, Lord E, Veronneau PY, Masonbrink R, Yu Q, Akker SED. The draft genome of Ditylenchus dipsaci. J Nematol. 2019;51:1-3. Epub 2019/05/28. doi: 10.21307/jofnem-2019-027. PubMed PMID: 31132003.

52.	Cotton JA, Lilley CJ, Jones LM, Kikuchi T, Reid AJ, Thorpe P, et al. The genome and life-stage specific transcriptomes of Globodera pallida elucidate key aspects of plant parasitism by a cyst nematode. Genome Biology. 2014;15(3). doi: ARTN R43

10.1186/gb-2014-15-3-r43. PubMed PMID: WOS:000338981300004.

53.	Eves-van den Akker S, Laetsch DR, Thorpe P, Lilley CJ, Danchin EGJ, Da Rocha M, et al. The genome of the yellow potato cyst nematode, Globodera rostochiensis, reveals insights into the basis of parasitism and virulence. Genome Biology. 2016;17. doi: ARTN 124

10.1186/s13059-016-0985-1. PubMed PMID: WOS:000378899700001.

54.	Masonbrink R, Maier TR, Muppirala U, Seetharam AS, Lord E, Juvale PS, et al. The genome of the soybean cyst nematode (Heterodera glycines) reveals complex patterns of duplications involved in the evolution of parasitism genes. Bmc Genomics. 2019;20. doi: ARTN 119

10.1186/s12864-019-5485-8. PubMed PMID: WOS:000458133500004.

55.	Sato K, Kadota Y, Gan P, Bino T, Uehara T, Yamaguchi K, et al. High-Quality Genome Sequence of the Root-Knot Nematode Meloidogyne arenaria Genotype A2-O. Microbiol Resour Ann. 2018;6(26). doi: UNSP e00519-18

10.1128/genomeA.00519-18. PubMed PMID: WOS:000452371900006.

56.	Blanc-Mathieu R, Perfus-Barbeoch L, Aury JM, Da Rocha M, Gouzy J, Sallet E, et al. Hybridization and polyploidy enable genomic plasticity without sex in the most devastating plant-parasitic nematodes. Plos Genetics. 2017;13(6). doi: ARTN e1006777

10.1371/journal.pgen.1006777. PubMed PMID: WOS:000404512600004.

57.	Szitenberg A, Salazar-Jaramillo L, Blok VC, Laetsch DR, Joseph S, Williamson VM, et al. Comparative Genomics of Apomictic Root-Knot Nematodes: Hybridization, Ploidy, and Dynamic Genome Change. Genome Biol Evol. 2017;9(10):2844-61. Epub 2017/10/17. doi: 10.1093/gbe/evx201. PubMed PMID: 29036290; PubMed Central PMCID: PMCPMC5737495.

58.	Somvanshi VS, Tathode M, Shukla RN, Rao U. Nematode Genome Announcement: A Draft Genome for Rice Root-Knot Nematode, Meloidogyne graminicola. J Nematol. 2018;50(2):111-6. doi: 10.21307/jofnem-2018-018. PubMed PMID: WOS:000455892800004.

59.	Opperman CH, Bird DM, Williamson VM, Rokhsar DS, Burke M, Cohn J, et al. Sequence and genetic map of Meloidogyne hapla: A compact nematode genome for plant parasitism. Proc Natl Acad Sci U S A. 2008;105(39):14802-7. Epub 2008/09/24. doi: 10.1073/pnas.0805946105. PubMed PMID: 18809916; PubMed Central PMCID: PMCPMC2547418.
